# Animal geolocation with convolution algorithms in Julia and R via Wahoo.jl

**DOI:** 10.1101/2025.09.19.677312

**Authors:** Edward Lavender, Carlo Albert, Andreas Scheidegger

**Affiliations:** Department of Systems Analysis, Integrated Assessment and Modelling, Eawag Swiss Federal Institute of Aquatic Science and Technology

**Keywords:** biologging, biotelemetry, filter, hidden Markov model, movement ecology, package, passive acoustic telemetry, state-space model

## Abstract

1. Animal geolocation is the core of movement ecology. In aquatic ecosystems, electronic tagging and tracking technologies, such as passive acoustic telemetry systems and biologging sensors, are widely deployed. However, statistical estimation of individual locations from these datasets can be challenging and computationally expensive.
2. Here, we introduce `Wahoo.jl`, a `Julia` package that fits state-space models to animal-tracking data via convolution algorithms. `Wahoo.jl` supports passive acoustic telemetry (detection/non-detection) and biologging (i.e., depth) datasets; implements grid-based filtering, smoothing and sampling of trajectories; and exploits GPU acceleration.
3. Using simulations, we illustrate how to use `Wahoo.jl` from `Julia` and `R` to reconstruct movements for an example individual tagged with an acoustic transmitter and an archival depth tag. We also provide validation and sensitivity analyses.
4. `Wahoo.jl` fills a key gap in the animal-tracking toolbox. The package provides an accessible, flexible and performant interface for an inference methodology that reliably handles multimodal inference problems that challenge other approaches. We discuss the approach’s pros and cons and provide guidance to readers on when to reach for `Wahoo.jl`.

## 1. Introduction

Animal geolocation is a core activity in ecology (Nathan et al., 2022). Studies of individual movements (Shaw, 2020), habitat selection (Florko et al., 2025) and conservation requirements (Hays et al., 2019) all require reliable estimates of individual locations. Electronic tagging and tracking technologies are widely deployed for this purpose (Hussey et al., 2015; Kays et al., 2015). In aquatic ecosystems, examples include satellite transmitters (which provide location fixes), passive acoustic telemetry systems (which record detections/non-detections of acoustic transmissions at static receivers) and biologging sensors (which record ancillary measurements, such as depths). Statistical animal geolocation leverages these data to infer individual locations (Jonsen et al., 2013; Lavender, Scheidegger, Moor, et al., 2025). This comprises two steps.

Step one is to formulate a state-space model for an individual’s locations (***s***) through time (*t*) (Jonsen et al., 2013). Here, we consider discrete time models (where *t* ∈ {1, …, *T*}). Generically, a discrete-time state-space model is a representation of hierarchical system with (i) a process model that describes how the system’s underlying state evolves, denoted *f*(***s***_1:*T*_), and (ii) an observation model, denoted *f*(***y***_1:*T*_ | ***s***_1:*T*_), that links the latent states to observations (***y***). If the latent states are discrete, the term hidden Markov model is often used (Thygesen et al., 2009). In a geolocation context, the process model is the movement process by which the individual’s state (location) unfolds and the observation model is a description of how observations (such as detections) arise contingent upon the state.

The second step is to infer the individual’s movement trajectories. Using Bayes Theorem, we can derive the joint probability of the states, given the observations, from the movement and observation processes as *f*(***s***_1:*T*_ | ***y***_1:*T*_) ∝ *f*(***s***_1:*T*_) *f*(***y***_1:*T*_ | ***s***_1:*T*_). To perform inference, we need to identify a suitable algorithm to approximate *f*(***s***_1:*T*_ | ***y***_1:*T*_) or related quantities (Lavender, Scheidegger, Moor, et al., 2025). Multiple inference approaches are available. Examples include filtering approaches (Thygesen et al., 2009), Laplace approximation (Jonsen et al., 2023) and Markov Chain Monte Carlo (MCMC) algorithms (Hostetter & Royle, 2020). For a review, see Lavender, Scheidegger, Moor, & Albert (2025).

Filtering algorithms are a natural choice for geolocation (Lavender, Scheidegger, Moor, et al., 2025). These algorithms consider the ‘partial’ marginal distribution *f*(***s***_*t*_ | ***y***_1:*t*_) of the individual’s state ***s***_*t*_ at time *t*, given all data ***y***_1:*t*_ up to that time. Filters exploit a recursive representation of *f*(***s***_*t*_ | ***y***_1:*t*_) in which the distribution *f*(***s***_*t*−1_ | ***y***_1:*t*−1_) is spread out by the movement process *f*(***s***_*t*_ | ***s***_*t*−1_) and then updated by the data *f*(***y***_*t*_ | ***s***_*t*_); i.e., *f*(***s***_*t*_ | ***y***_1:*t*_) ∝ ∫ *f*(***s***_*t*−1_ | ***y***_1:*t*−1_) *f*(***s***_*t*_ | ***s***_*t*−1_) *f*(***y***_*t*_ | ***s***_*t*_) *d****s***_*t*−1_. These two steps are termed the ‘prediction’ and ‘update’ (Thygesen et al., 2009). With an additional smoothing step, the full marginal *f*(***s***_*t*_ | ***y***_1:*T*_) can be computed. Trajectories can also be sampled from the joint distribution *f*(***s***_1:*T*_ | ***y***_1:*T*_). Inference focuses on the states (though it is possible to estimate static parameters in the movement and observation processes via the likelihood).

Filtering approaches include Kalman filters (Sibert et al., 2003), particle filters (Lavender, Scheidegger, Albert, et al., 2025a) and grid-based filters (Thygesen et al., 2009). Kalman filtering is an efficient option in linear systems with Gaussian errors (Sibert et al., 2003), but not directly applicable in passive acoustic telemetry systems (with binary observations). Particle filtering is a more general approach (Doucet & Johansen, 2009). Particle filters approximate *f*(***s***_*t*_ | ***y***_1:*t*_) using weighted samples termed particles. Given a sample of particles for time *t* − 1, particle filters recursively sample new particles for *t* from the movement model, weight them via the observation model and then resample them accordingly. The `patter` and `Patter.jl` packages implement this approach (Lavender, Scheidegger, Albert, et al., 2025b). While powerful, particle filters can be affected by particle degeneracy—the concentration of weights around a few particles, which can lead to poor approximations of complex probability distributions (Doucet & Johansen, 2009). Extreme particle degeneracy was observed in a study integrating acoustic and depth observations from flapper skate (*Dipturus intermedius*) in a bathymetrically complex environment (Lavender, Scheidegger, Albert, et al., 2025). Inference in this context is akin to finding a route through a bathymetric maze, which is a hard task for sampling methods (since particles simulated from the movement process are typically incompatible with the observations).

Grid-based filtering is a solution (Thygesen et al., 2009). In a grid-based filter, we discretise distribution *f*(***s***_*t*_ | ***y***_1:*t*_) over the study area. Like a particle filter, the distribution is spread out in every time step (the prediction) and then corrected by observations (the update). However, unlike a particle filter, probabilities are computed for each cell directly, so particle degeneracy is not an issue. Grid-based filters were introduced by Pedersen et al. (2008) and Thygesen et al. (2009) for tidal geolocation of demersal fish. Subsequently, the approach was adapted for light-level and oceanographic data collected by pop-up satellite archival transmitters (Braun et al., 2018). By avoiding sampling, the approach reliably represents multimodal distributions but can be computationally expensive. However, most computations can be parallelised and, for certain kinds of movement models, the prediction can be implemented as a two-dimensional convolution, for which optimised GPU routines are available.

Here, we provide a performant `Julia` implementation of grid-based filtering for geolocation. `Julia` is a compiled programming language with strong GPU support that maintains a high-level experience (Bezanson et al., 2017). Our package is designed for passive acoustic telemetry and biologging datasets (complementing the `patter` packages). We have implemented filtering, smoothing and sampling of trajectories and exploit convolution algorithms to alleviate the computational challenge of gridding. The package enables the acoustic telemetry community to exploit grid-based filtering and should support reliable geolocation analyses in many systems. In what follows, we formalise the geolocation model and the inference approach. We provide worked examples in `Julia` and `R` plus validation and sensitivity analyses.

## 2. Methodology

This section outlines a Bayesian state-space model for the location of a tagged animal (following Lavender et al., 2025a) and the inference procedure (following Thygesen et al., 2009). For full details, see Supporting Information §1. Table S1 provides a mapping between our and Thygesen et al.’s (2009) notation.

### 2.1 Model formulation

#### Posterior

Our goal is to estimate a tagged animal’s location through time, accounting for the movement process and the observations. The individual’s unknown trajectory is represented probabilistically by the joint distribution *f*(***s***_1:*T*_ | ***y***_1:*T*_) of the individual’s locations (***s*** *=* (*s*_*x*_, *s*_*y*_)) and the observations (***y***); that is,

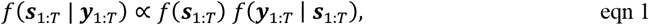

where *f*(***s***_1:*T*_) and *f*(***y***_1:*T*_ | ***s***_1:*T*_) denote the movement and observation processes, respectively.

#### Movement

We model *f*(***s***_1:*T*_) with a probability distribution *f*(***s***_*t=*1_) for the individual’s initial location and a Gaussian random walk *f*(***s***_*t*_ | ***s***_*t*−1_) for the movement process from ***s***_*t*−1_ → ***s***_*t*_, i.e.,

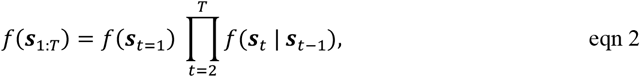

Where

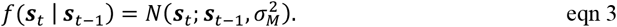

#### Observations

We model the observation process *f*(***y***_1:*T*_ | ***s***_1:*T*_) as

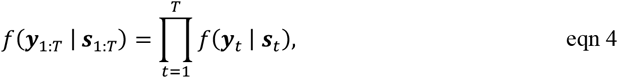

assuming independence. The term *f*(***y***_*t*_ | ***s***_*t*_) is data-specific (Lavender, Scheidegger, Albert, et al., 2025a). For example, for acoustic observations, a Bernoulli model in which the probability of a detection declines with distance from a receiver is appropriate. Other observations, such as depth measurements (together with knowledge of the bathymetry), can also be included in this term (see Supporting Information §1.1).

### 2.2. Inference

Inference comprises grid-based filtering, smoothing and sampling (see Supporting Information §1.2). The filter targets *f*(***s***_*t*_ | ***y***_1:*t*_). An approximation of *f*(***s***_*t*_ | ***y***_1:*t*_) is achieved by discretisation. We represent the study area as a matrix and recursively estimate the probability *P*_*i,j,t*_(***y***_1:*t*_) of the individual being in each grid cell *S*_*i*_,_*j*_ with coordinates (*i, j*) at time *t*, conditional on the observations ***y***_1:*t*_ (see Supporting Information §1.2.1). At each time step, the computation is a two-stage process: first, the movement process that diffuses the probability distribution is computed via convolution; then the probabilities are weighted based on the observations. The full marginal probabilities *P*_*i,j,t*_(***y***_1:*T*_) are computed via smoothing (see Supporting Information §1.2.2). The overall occupancy distribution is defined across the grid by the normalised sum of all *P*_*i,j,t*_(***y***_1:*T*_) values. Trajectories are sampled by a recursive algorithm that conditions on sampled positions and observations (see Supporting Information §1.2.3). Static parameters can be estimated over multiple filter runs via the likelihood (see Supporting Information §1.2.4).

## 3. Implementation

Our package `Wahoo.jl` implements grid-based filtering, smoothing and sampling of trajectories (Scheidegger, 2025). The package name draws inspiration from the wahoo (*Acanthocybium solandri*), a fast-swimming mackerel species. We designed the package to be lightweight, low-dependency and performant.

`Wahoo.jl` focuses on inference and exports a single function, `track()`, for this purpose. To implement the function, users should provide a bathymetry raster (`GeoArray`) for the study area and a probability distribution for the individual’s initial location. The movement process (eqn 3) is implemented as a two-dimensional convolution, representing a random walk with a user-defined standard deviation. Observations are provided as a `Vector`, with one element for each sensor (e.g., receiver) that contains the observation time series. Accompanying `Vector`s with `Function`s defining the observational models (*f*(***y***_*t*_ | ***s***_*t*_) in eqn 4) and sensor positions (if applicable) should also be provided. We provide example models; custom functions are also supported. Using these inputs, `track()` performs inference using the CPU or `CUDA`-compatible GPUs.

Outputs include filtered and/or smoothed probability distributions, the occupancy distribution, trajectories and a log-probability vector. To handle memory requirements, distributions are recorded at a subset of time (`tsave`) steps. Outputs are defined in `Array`s or `Vector`s, which can be analysed using standard routines in any programming language. Example analyses include analyses of centres of activity, occupancy, home ranges and habitat selection (Lavender, Scheidegger, Moor, et al., 2025).

For estimation of static parameters, users can plug the log-likelihood estimate into a grid-search or optimisation algorithm, or perform Bayesian inference using any dedicated `Julia` package.

## 4. Examples

### 4.1. Workflow

We provide an example workflow using simulation (Fig. 1). The example is motivated by previous skate research (Lavender et al., 2021b, 2021a; Lavender, Scheidegger, Albert, et al., 2025). On a 100 x 100 m bathymetry raster (557,568 cells), we simulated a movement trajectory at a time resolution of two minutes over a one month period (22,320 time steps), plus acoustic and depth observations for a hypothetical skate, following Fig. 1 (see Supporting Information §2.1 for the mathematics). Steps for inference with `Wahoo.jl` are below. The supporting materials provide complete workflows in `Julia` and `R`.

**Fig. 1.**
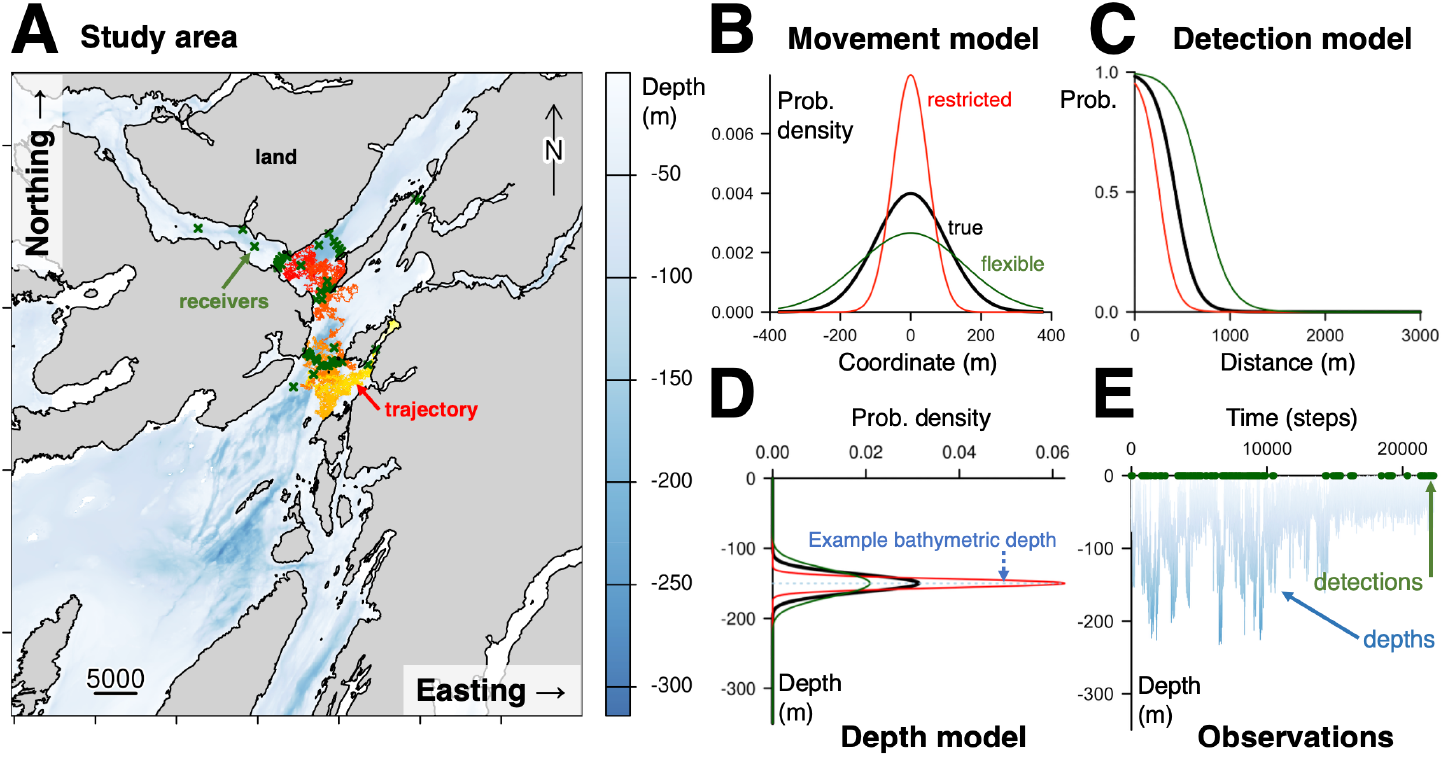
An example state-space model for animal geolocation. **A** shows a study area, including a simulated trajectory, coloured by time (yellow to red), and acoustic receivers. **B–D** depict model components: a Gaussian movement model; a Bernoulli acoustic observation model in which detection probability declines with distance from a receiver; and a truncated Gaussian depth observation model centred on the seabed (skate are predominantly benthic). In the example workflow, we simulated and analysed data using the same processes (black). In sensitivity analyses, we reimplemented algorithms with restricted (red) and flexible (green) model parameterisations. **E** shows simulated observations.

#### A. Load packages

First, we load necessary packages.

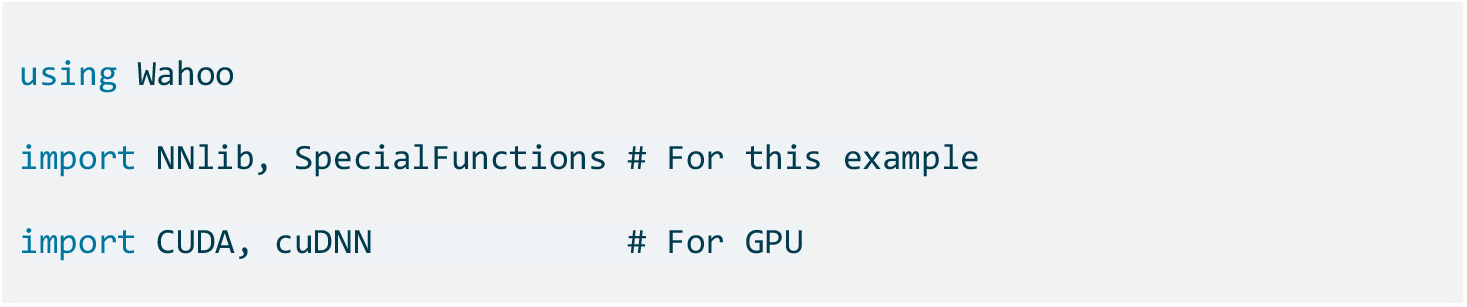

#### B. Define inputs

Second, we define inputs, including a bathymetry `GeoArray` (`bathy`), a time stamp `Vector` for the study period (`timeline`), receiver positions (`acoustic_pos`), a `Vector` of acoustic observation time series at each of 48 receivers (`acoustic_obs`) and a depth `Vector` (`depth_obs`). We also define a uniform distribution (`Matrix`) for the individual’s initial position (`p0`).

#### C. Formulate state-space model

Third, we formulate our model (Fig. 1) by defining the movement standard deviation and observation functions that evaluate 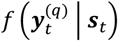 for each data type (*q*).

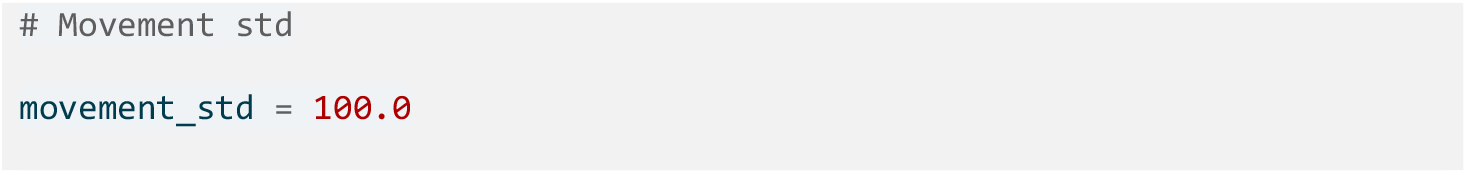

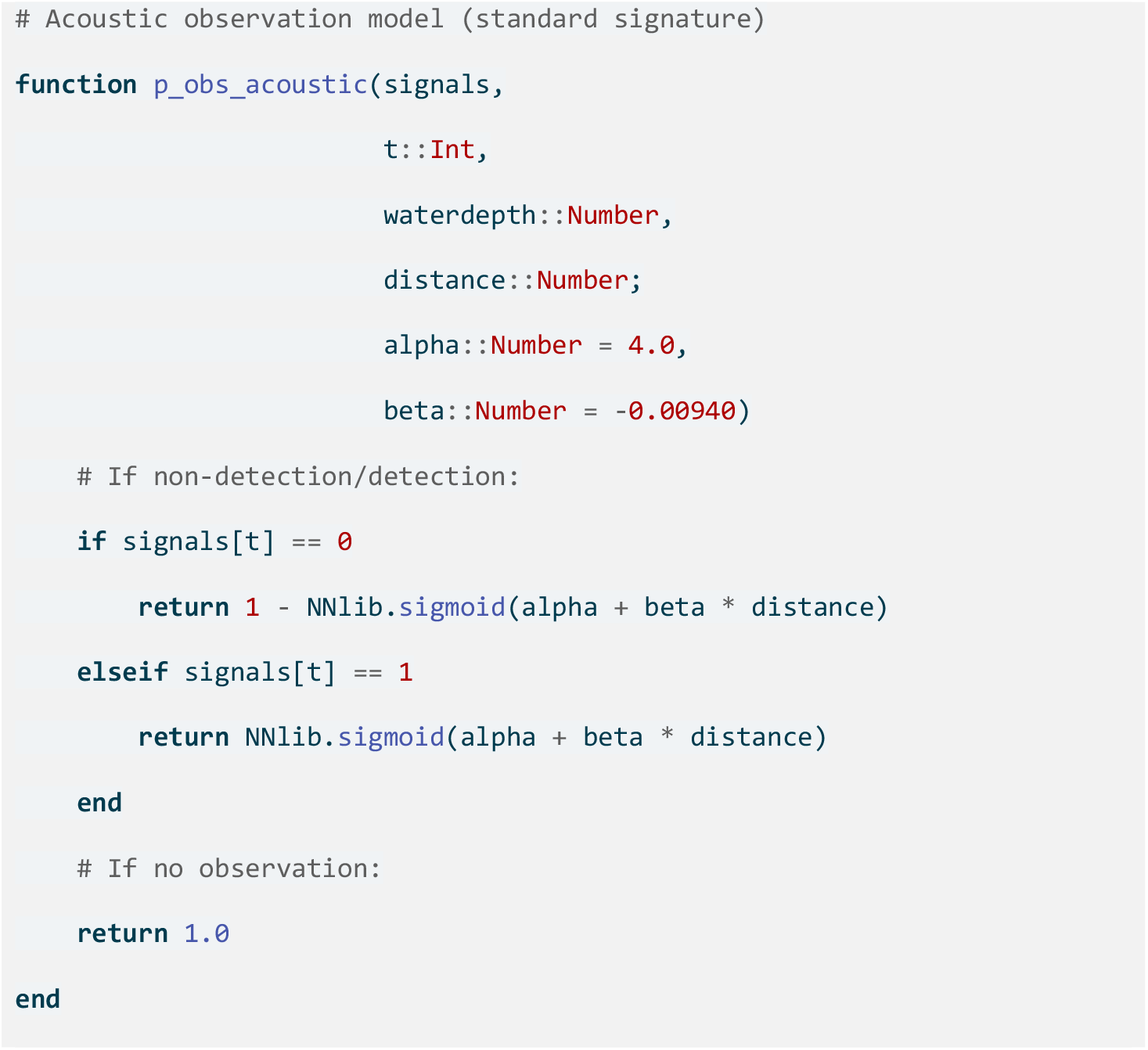

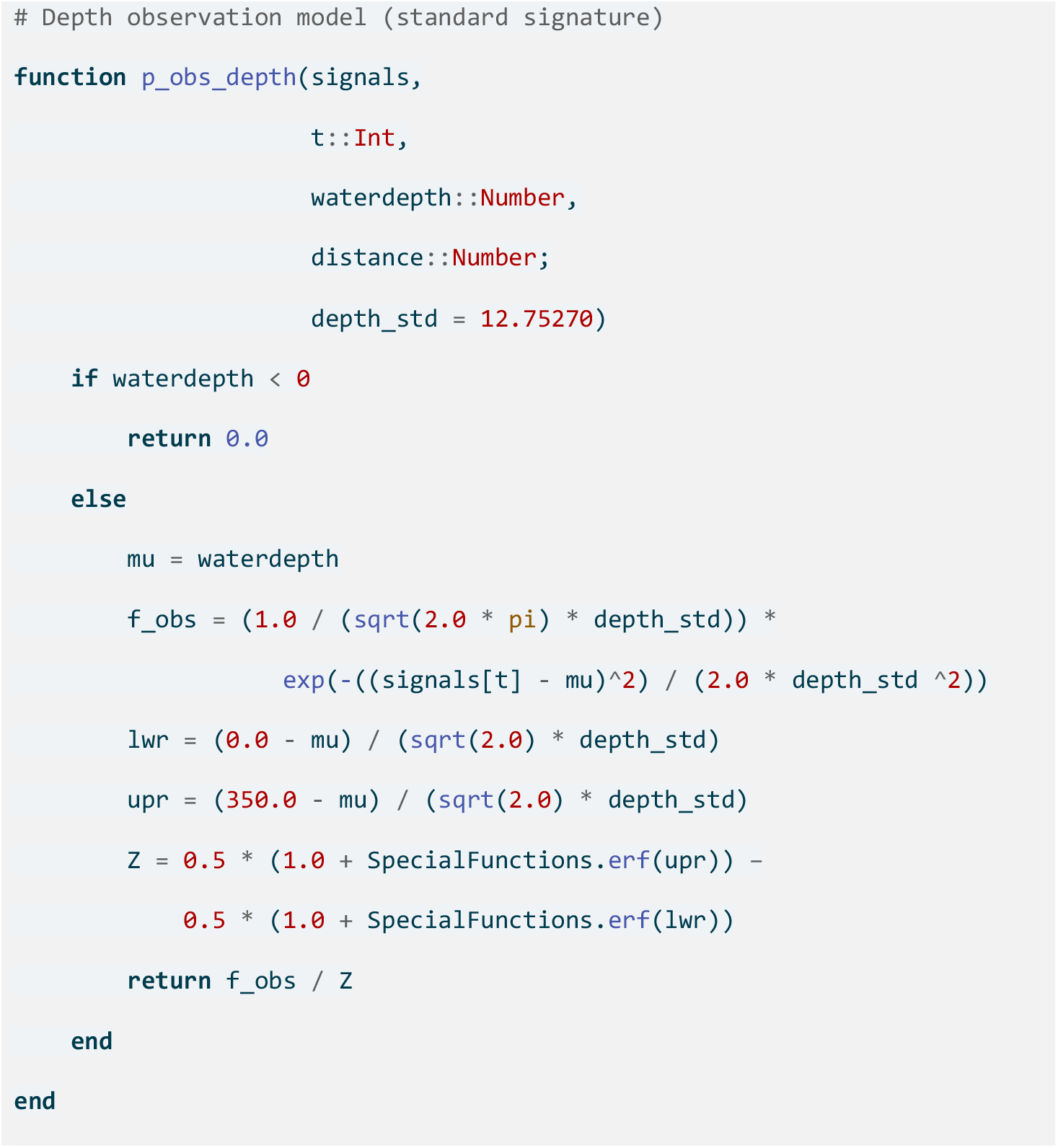

#### D. Run inference

Finally, we perform inference. For the filter and smoother, we save every 60^th^ distribution.

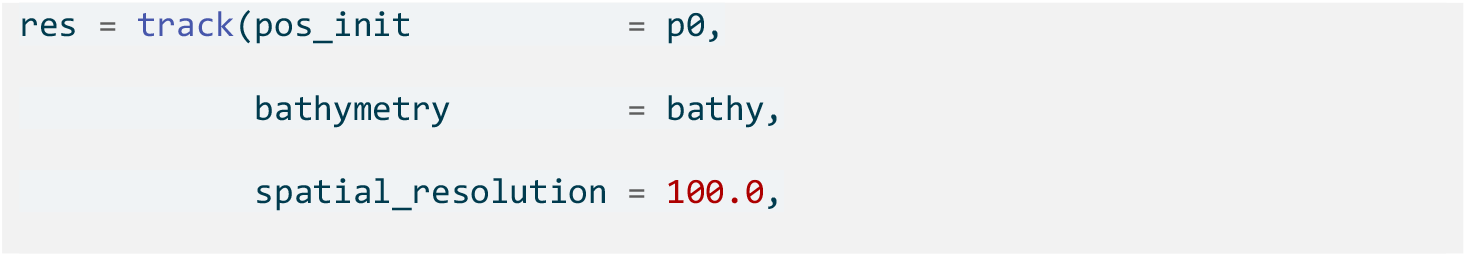

Hidden Markov models for geolocation

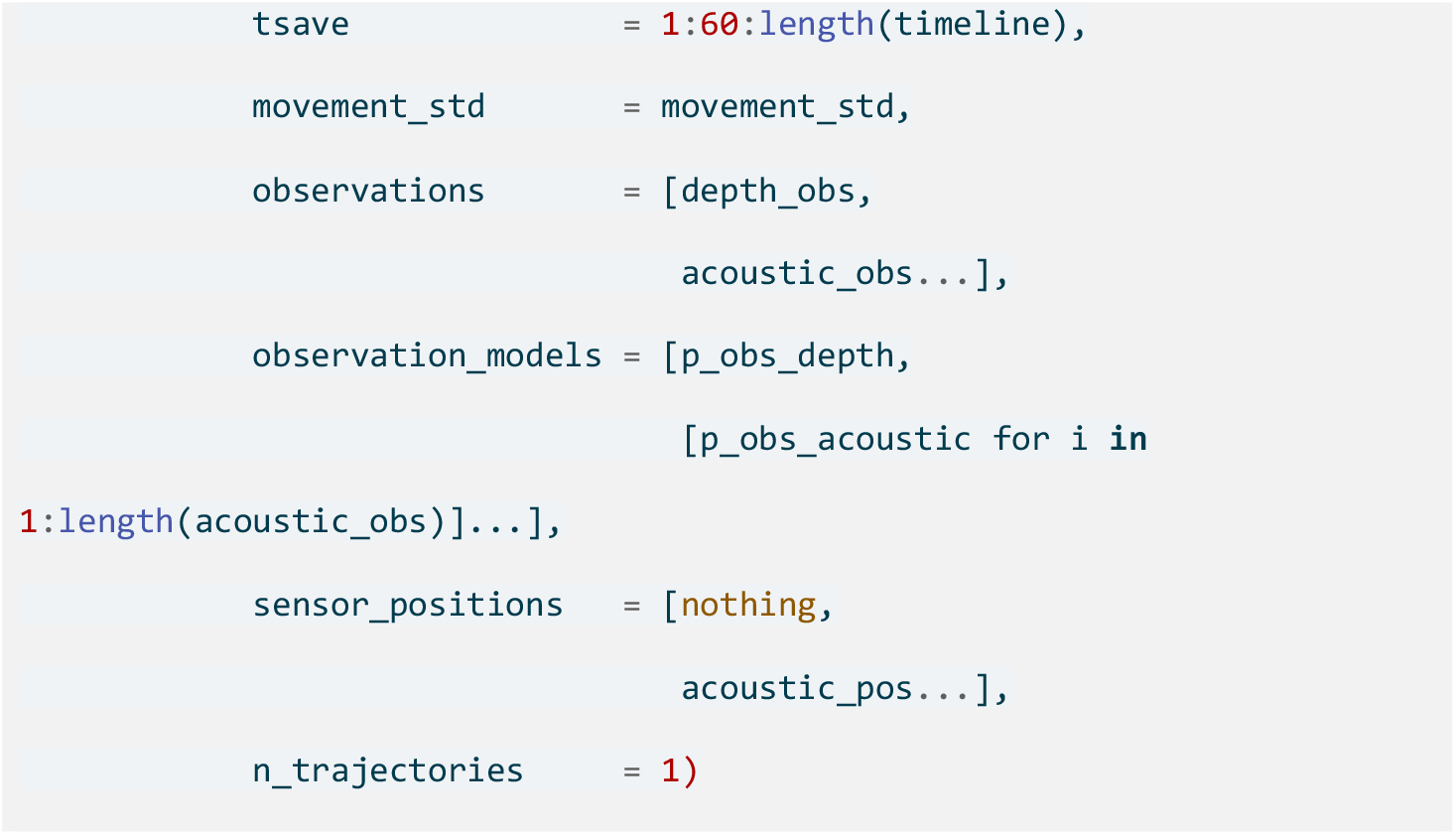

The output is a `NamedTuple` that we can analyse as desired. Here, we visualise simulated and inferred movements.

### 4.2. Validation and sensitivity

To further illustrate `Wahoo.jl`, we repeated the simulation for five hypothetical individuals. As a validation exercise, we reconstructed movements using data-generating parameters with both `Wahoo.jl` and `patter` (see Supporting Information §2.2). We also illustrate a sensitivity analysis that evaluates epistemic uncertainty by repeating our analysis with mis-specified parameters (see Supporting Information §2.3 and Table S2 for parameter settings). Code is available online (Lavender, Scheidegger, & Albert, 2025).

## 5. Results

In the example analysis, we inferred the individual’s locations across a 557,568-cell grid over 22,320 time steps. Simulated and inferred occupancy distributions matched closely (Fig. 2). Inference, including filtering (6 min), smoothing (23 min) and sampling one trajectory (23 min), took approximately one hour on a 2018 consumer-grade NVIDIA GeForce RTX 2080 Ti GPU. On the CPU (2019 AMD EPYC 7742 server hardware), computation time was 3.0 hours (filter), 4.0 hours (smoother) and 3.9 hours (sampler). For comparison, a preliminary analysis with `patter` using 20,000/1,000 filter/smoother particles took approximately 1 hour (single-threaded on the CPU). The full analysis (100,000/2,000 particles) took 8.6 hours.

**Fig. 2.**
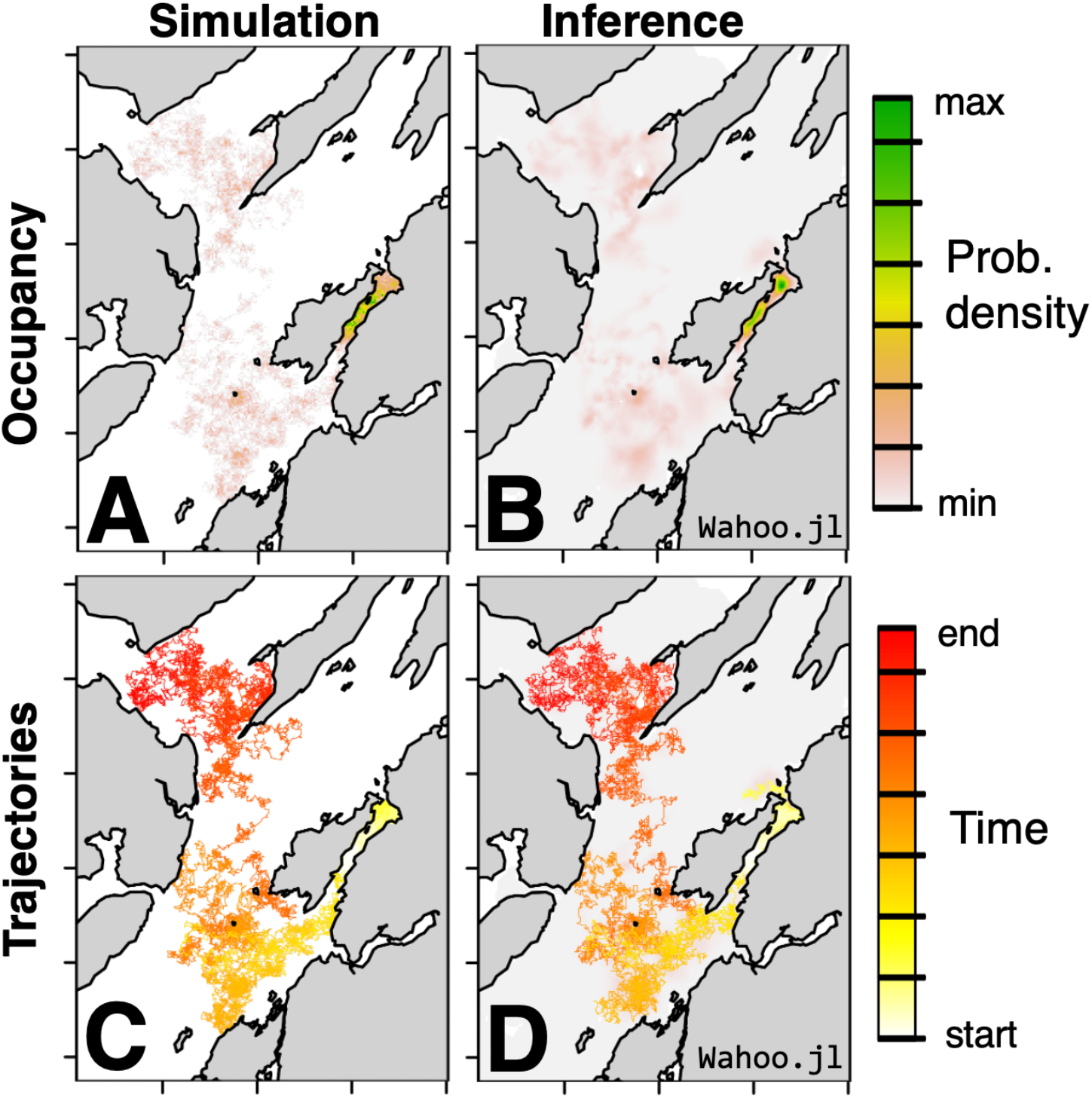
A comparison of simulated and inferred patterns for an example individual. **A**–**B** show occupancy distributions; **C–D** show the simulated trajectory versus a sampled trajectory.

In the validation analysis, occupancy distributions reconstructed by `Wahoo.jl` and `patter` broadly aligned, providing high-level validation of the routines (Fig. S1). However, effective particle sample sizes were relatively low (indicating degeneracy). Particle algorithms were thus associated with somewhat patchier distributions. This effect was exaggerated in the simulation with the lowest sampling efficacy.

In the sensitivity analysis, occupancy distributions were broadly robust to parameter misspecification (Fig. S2). Additionally, parameter misspecification appeared to degrade the numerical stability of `patter`, unlike `Wahoo.jl` (Fig. S3).

## 6. Discussion

`Wahoo.jl` provides an accessible, flexible and efficient implementation of convolution algorithms for geolocation (Thygesen et al., 2009). The inference methodology reliably handles geolocation problems in complex landscapes while exhibiting predictable computational performance. As a case study, we are deploying `Wahoo.jl` in a substantial analysis of our skate datasets. We hope `Wahoo.jl` will support integrative geolocation analyses in many other systems.

To apply `Wahoo.jl`, users should formulate a state-space model for inference. The movement model implemented by `Wahoo.jl` is a Gaussian random walk, with a customisable standard deviation. For this model, the probability density of a movement depends only on the distance between states, which enables efficient implementation via convolution. From a Bayesian perspective, we also believe this is a widely applicable model for our belief about *f*(***s***_*t*_ | ***s***_*t*−1_). The observation model design is flexible. We encourage users to leverage data, domain expertise and literature for model development (Lavender, Scheidegger, Albert, et al., 2025a, 2025b). In acoustic telemetry systems, observations are typically sparse and we prefer to focus inference on the latent states. Users seeking to estimate static parameters should evaluate data sufficiency and computational requirements via simulation.

`Wahoo.jl` is one of the few model-based acoustic-telemetry packages suitable for direct use by practitioners (Lavender, Scheidegger, Moor, et al., 2025). The key strength of the inference procedure is the use of spatial discretisation, which avoids the convergence issues that can affect particle filters. As a result, complex multimodal distributions can be inferred reliably. Unlike simulation approaches, computation time is also predictable. `Wahoo.jl` supports applications on both CPUs and GPUs (which is faster). Computation time is competitive with `patter`, even with half a million grid cells. Comparison with grid-based routines is difficult in the absence of standardisation, but it is noteworthy to estimate individual locations across more than 500,000 cells on over 20,000 time steps in less than one hour (*cf*. Pedersen et al., 2008). This `Julia` implementation with GPU-acceleration should be highly competitive.

Limitations of the inference methodology implemented by `Wahoo.jl` include restrictions on the movement model and computational requirements (notwithstanding GPU-acceleration). While filtering is fast, smoothing, sampling trajectories and estimating static parameters are comparably expensive. Memory demand (especially on the GPU) is another challenge (for long time series and large grids). For alternative approaches that make different trade-offs, we direct readers to the following review (Lavender, Scheidegger, Moor, et al., 2025).

For `Wahoo.jl`, community feedback will support package development. Research avenues include continued optimisation, enhanced flexibility (with support for more data and GPU types) and the development of wrappers for other languages. More generally, there is a need to benchmark available approaches to understand how problem complexity, algorithmic design and software implementations shape the suitability of different approaches in different settings (Lavender et al., 2023; Lavender, Scheidegger, Moor, et al., 2025).

## Supporting information

Supporting Code (Julia)

## Author contributions

All authors conceived the project. Andreas Scheidegger developed the `Wahoo.jl` package. Carlo Albert developed the mathematical notation. Edward Lavender developed the manuscript. All authors shared expertise, provided inputs and approved publication.

## Data availability statement

The `Wahoo.jl` package is available on GitHub (Scheidegger, 2025). Data and code for this study are archived on Zenodo (Lavender, Scheidegger, & Albert, 2025).

## Acknowledgments

We are grateful to Helen Moor for supporting this project.

