## Supporting Information for "Animal geolocation with convolution algorithms in Julia and R via Wahoo.jl"

**Contents** **Pages**


| | | |
|---|---|---|
| **1.** | **Methodology** | **1–8** |
| 1.1. | Model formulation | 1–3 |
| 1.1.1. | Posterior | 1 |
| 1.1.2. | Prior | 1–2 |
| 1.1.3. | Likelihood | 2–3 |
| 1.2. | Inference | 3–8 |
| 1.2.1. | Filtering | 4–6 |
| 1.2.2. | Smoothing | 6–7 |
| 1.2.3. | Sampling trajectories | 7–8 |
| 1.2.4. | Parameter estimation via the likelihood | 8 |
| **2.** | **Simulation analyses** | **8–11** |
| 2.1. | Example workflow | 8–10 |
| 2.2. | Validation analyses | 10–11 |
| 2.3. | Sensitivity analyses | 11 |
| | **References** | **12–13** |


For example workflows written in `Julia` and `R`, see the supporting documents.

For supporting figures, see the supporting figures document.

For supporting tables, see the supporting tables document.

### 1. Methodology

In the Main Text, we outline a generic state-space model for animal geolocation (following Lavender et al., 2025a) and a filtering algorithm (Thygesen et al., 2009). This Supporting Information provides full details. We first recap and exemplify the model formulation (§1.1) before formalising the inference procedure (§1.2).

#### 1.1. Model formulation

##### 1.1.1. Posterior

We consider a state-space model for joint distribution $f(\boldsymbol{s}_{1:T} \mid \boldsymbol{y}_{1:T})$ of an individual's trajectories $\boldsymbol{s}_{1:T}$ conditional on the observations $\boldsymbol{y}_{1:T}$ through time ($t \in \{1, \dots, T\}$). By Bayes' Theorem, this distribution is proportional to the product of a prior and the likelihood, that is,

$$f(\boldsymbol{s}_{1:T} \mid \boldsymbol{y}_{1:T}) \propto f(\boldsymbol{s}_{1:T})\, f(\boldsymbol{y}_{1:T} \mid \boldsymbol{s}_{1:T}), \qquad \text{eqn 1}$$

where $f(\boldsymbol{s}_{1:T})$ denotes the prior and $f(\boldsymbol{y}_{1:T} \mid \boldsymbol{s}_{1:T})$ denotes the likelihood.

##### 1.1.2. Prior

The prior represents the movement process. We model the prior with a probability density distribution for the individual's initial location, $f(\boldsymbol{s}_{t=1})$, and a Markovian movement model, $f(\boldsymbol{s}_{t+1} \mid \boldsymbol{s}_t)$, that describes the probability density of movement between two locations, $\boldsymbol{s}_t \rightarrow \boldsymbol{s}_{t+1}$:

$$f(\boldsymbol{s}_{1:T}) = f(\boldsymbol{s}_{t=1}) \prod_{t=1}^{T-1} f(\boldsymbol{s}_{t+1} \mid \boldsymbol{s}_t). \qquad \text{eqn 2}$$

As a model for $f(\boldsymbol{s}_{t+1} \mid \boldsymbol{s}_t)$, we consider a Gaussian random walk in which the predicted distribution of an individual's location at time $t+1$ is centred on the location at time $t$, with a standard deviation $\sigma_M$ related to the individual's movement behaviour:

$$f(\boldsymbol{s}_{t+1} \mid \boldsymbol{s}_t) = N(\boldsymbol{s}_{t+1}; \boldsymbol{s}_t, \sigma_M^2). \qquad \text{eqn 3}$$

This is a continuous-space, discrete-time representation of a diffusion equation (excluding drift)[1]. The standard deviation of a diffusion process is related to a diffusivity parameter $D$ via $\sigma_M = \sqrt{2D\Delta t}$, where $\Delta t$ is the time step duration.

##### 1.1.3. Likelihood

The term $f(\boldsymbol{y}_{1:T} \mid \boldsymbol{s}_{1:T})$ in eqn 1 denotes the joint likelihood. This term measures the probability of the observations given the individual's locations $\boldsymbol{s}_{1:T}$. We model $f(\boldsymbol{y}_{1:T} \mid \boldsymbol{s}_{1:T})$ as the product of the likelihood from each time step,

$$f(\boldsymbol{y}_{1:T} \mid \boldsymbol{s}_{1:T}) = \prod_{t=1}^{T} f(\boldsymbol{y}_t \mid \boldsymbol{s}_t), \qquad \text{eqn 4}$$

assuming independence[2].

The term $f(\boldsymbol{y}_t \mid \boldsymbol{s}_t)$ can be formulated for different data types and customised in line with study-specific considerations (Lavender, Scheidegger, Albert, Biber, Illian, et al., 2025a). In our worked examples (see Main Text), we consider acoustic $\boldsymbol{y}_t^{(A)}$ and archival depth $y_t^{(D)}$ observations from flapper skate (*Dipturus intermedius*) and the combined likelihood

$$f(\boldsymbol{y}_t \mid \boldsymbol{s}_t) = f\left(\boldsymbol{y}_t^{(A)} \,\middle|\, \boldsymbol{s}_t\right) f\left(y_t^{(D)} \,\middle|\, \boldsymbol{s}_t\right). \qquad \text{eqn 5}$$

**Acoustic observations.** Acoustic observations $\boldsymbol{y}_t^{(A)}$ comprise detections (1) and non-detections (0) at each operational receiver $k$ (that is, $y_{t,k}^{(A)} \in \{0, 1\}$). We modelled the acoustic observation process using a Bernoulli model

$$f\left(y_{t,k}^{(A)} \,\middle|\, \boldsymbol{s}_t\right) = \text{Bernoulli}\left(p_{t,k}(\boldsymbol{s}_t)\right) \qquad \text{eqn 6}$$

---

[1] During model inference, filtering algorithms typically focus on estimation of the latent states while assuming static parameters, such as $\sigma_M^2$, in the movement and observation models are known (see §1.2). In our worked examples, we set static parameters based on prior research (Lavender, Scheidegger, Albert, Biber, Aleynik, et al., 2025) and illustrate a sensitivity analysis that evaluates epistemic uncertainty (see §2.3). The parameter $\sigma_M$ is called `movement_std` in the code. However, with sufficient data and computational resources, joint estimation of latent states and static parameters is possible (see §1.2.4).

[2] By updating our prior $f(\boldsymbol{s}_{1:T})$ for the individual's location with the likelihood, this model represents the distribution of possible locations for an individual as (a) those locations which can be reached which are also (b) compatible with the observations.

in which the probability of detection $p_{t,k}$ at time $t$ at receiver $k$ declines logistically with the distance between the transmitter (in location $\boldsymbol{s}_t$) and the receiver (in location $\boldsymbol{r}_k$)

$$p_{t,k}(\boldsymbol{s}_t) = \left(1 + e^{-(\alpha - \beta \times |\boldsymbol{s}_t - \boldsymbol{r}_k|)}\right)^{-1}, \qquad \text{eqn 7}$$

in line with the coefficients $\alpha$ and $\beta$[3]. Assuming independence, the combined likelihood across all receivers is

$$f\left(\boldsymbol{y}_t^{(A)} \mid \boldsymbol{s}_t\right) = \prod_k f\left(y_{t,k}^{(A)} \mid \boldsymbol{s}_t\right), \qquad \text{eqn 8}$$

where $k$ indexes over operational receivers at time $t$.

**Depth observations.** Archival depth observations $y_t^{(D)}$ comprise animal-borne depth (m) measurements. In our worked examples, we model depth observations for (simulated) flapper skate, which are considered predominately benthic animals. For this species, we formulate a Gaussian model

$$f\left(y_t^{(D)} \mid \boldsymbol{s}_t\right) = \text{truncated}(N(b(\boldsymbol{s}_t), \sigma_D^2), 0, 350), \qquad \text{eqn 9}$$

in which the individual's depth is assumed to be normally distributed around the bathymetric depth $b(\boldsymbol{s}_t)$ with variance $\sigma_D^2$. The $\sigma_D$ parameter captures uncertainty in the bathymetric and observed depth measurements as well as the degree to which flapper skate are considered benthic animals[4] (see §2.1). We truncated the distribution between the surface and the maximum depth in the study area. In other systems, models should be customised in line with species' biology and system design.

#### 1.2. Inference

We have formulated an example model for the joint distribution $f(\boldsymbol{s}_{1:T} \mid \boldsymbol{y}_{1:T})$ of an individual's locations, given the observations. We now outline the grid-based inference algorithm.

---

[3] In the supporting code, these parameters are coded as `receiver_alpha` and `receiver_beta`. In previous work with particle algorithms (Lavender, Scheidegger, Albert, Biber, Illian, et al., 2025a, 2025b), we used a similar model truncated by a maximum detection range parameter. The truncation can alleviate particle degeneracy in the particle algorithms via acoustic containerisation. Grid-based filtering avoids particle degeneracy, so for the purpose of this manuscript we consider the simpler (untruncated) model.

[4] In the example code, $b(\boldsymbol{s}_t)$ is coded as `mu = waterdepth`. The $\sigma_D$ parameter is coded as `depth_std`.

##### 1.2.1. Filtering

We first consider the 'partial' marginal distribution $f(\boldsymbol{s}_{t+1} \mid \boldsymbol{y}_{1:t+1})$ of the individual's location at a given time step, given the data up to and including that time. This distribution can be represented in terms of the movement process $f(\boldsymbol{s}_{t+1} \mid \boldsymbol{s}_t)$ and the likelihood $f(\boldsymbol{y}_{t+1} \mid \boldsymbol{s}_{t+1})$:

$$f(\boldsymbol{s}_{t+1} \mid \boldsymbol{y}_{1:t+1}) = \frac{1}{Z(\boldsymbol{y}_{1:t+1})} \int f(\boldsymbol{y}_{t+1} \mid \boldsymbol{s}_{t+1})\, f(\boldsymbol{s}_{t+1} \mid \boldsymbol{s}_t)\, f(\boldsymbol{s}_t \mid \boldsymbol{y}_{1:t})\; d\boldsymbol{s}_t$$
$$= \frac{1}{Z(\boldsymbol{y}_{1:t+1})}\, f(\boldsymbol{y}_{t+1} \mid \boldsymbol{s}_{t+1})\, f(\boldsymbol{s}_{t+1} \mid \boldsymbol{y}_{1:t}), \qquad \text{eqn 10}$$

where $Z(\boldsymbol{y}_{1:t+1}) = \int d\boldsymbol{s}_{t+1}\, f(\boldsymbol{y}_{t+1} \mid \boldsymbol{s}_{t+1})\, f(\boldsymbol{s}_{t+1} \mid \boldsymbol{y}_{1:t})$ is the normalisation constant.

The forward filter achieves an approximation of $f(\boldsymbol{s}_{t+1} \mid \boldsymbol{y}_{1:t+1})$ by discretisation. The general theory, in discretised form, is as follows. We represent the study area (the state-space of possible locations) as a matrix. We compute the probability $P_{ij,t}(\boldsymbol{y}_{1:\tau})$ of the individual being in grid cell $S_{i,j}$ with coordinates $(i, j)$ at time $t$, conditional on the data from time $1{:}\tau$, where τ is a generic placeholder for the time step (such as $t$, $t+1$ and $T$). In the forward filter, estimates of $P_{ij,t}(\boldsymbol{y}_{1:t})$ are computed recursively by an algorithm that steps forwards in time in three stages. Starting with an initial probability distribution for the location of the animal (**A**), we iteratively diffuse the distribution via convolution, in line with the animal's movement behaviour (**B**), before updating the resulting probabilities in each cell in line with their compatibility with the data (**C**). This works as follows.

**A. Initialisation.** We initialise a probability distribution for the individual's initial location on the grid, $P_{ij,t=1}(\boldsymbol{y}_{t=1})$. If the initial position is known, this distribution is simply one in the corresponding grid cell and zero elsewhere.

**B. Movement.** Second, we predict the individual's distribution for the next time step, by updating the distribution for the current time step with transition (movement) probabilities derived from the movement model (eqn 3). That is, we predict the probability $P_{ij,t+1}(\boldsymbol{y}_{1:t})$ of the individual being in each grid cell, accounting for all possible moves into that cell (from $S_{i'j'} \rightarrow S_{ij}$) and the previous data:

$$P_{ij,t+1}(\boldsymbol{y}_{1:t}) = \sum_{i'j'} P_{ij \mid i'j'}\, P_{i'j',t}(\boldsymbol{y}_{1:t}), \qquad \text{eqn 11}$$

where $P_{ij\,|\,i'j'}$ represents the probability of the move from $S_{i'j'} \rightarrow S_{ij}$. With the assumption of a Gaussian random walk and by choosing the time step duration to be sufficiently small such that an individual can only move between neighbouring grid cells over the time from $t \rightarrow t+1$, we can efficiently approximate $P_{ij,t+1}(\boldsymbol{y}_{1:t})$ via convolution. Let $H_{rs}$ denote a 3-by-3 transition (movement) matrix $H_{rs}$ that encodes the probability of moving into a neighbouring cell, with coordinates $(i+r, j+s)$, from the current cell, at $(i,j)$, so $r$ and $s$ run over $\{0, \pm 1\}$. This allows us to express eqn 11 as a convolution operation and slide $H_{rs}$ over the grid to update the probability of the individual being in each cell at time $t$:

$$P_{ij,t+1}(\boldsymbol{y}_{1:t}) = \sum_{r,s\,\in\,\{0,\pm 1\}} H_{rs}\, P_{i+r,j+s,t}(\boldsymbol{y}_{1:t}). \qquad \text{eqn 12}$$

The elements of $H_{rs}$ are linked to the individual's movement behaviour via the diffusivity parameter, as shown by finite difference discretisation of the movement (diffusion) kernel, whereby:

$$H_{rs} = \begin{bmatrix} 0 & 0 & 0 \\ 0 & 1 & 0 \\ 0 & 0 & 0 \end{bmatrix} + \frac{D\Delta \mathrm{t}}{h^2} \begin{bmatrix} 0 & 1 & 0 \\ 1 & -4 & 1 \\ 0 & 1 & 0 \end{bmatrix}, \qquad \text{eqn 13}$$

where $D = \sigma_M^2 / 2\Delta t$ is the diffusivity (which can be interpreted in terms of the variance of a random walk), $h$ is the grid resolution and $\Delta t$ is the time step duration (Versteeg & Malalasekera, 1995). To ensure that transition probabilities $H_{rs}$ are non-negative, the spatial ($h$) and temporal ($\Delta \mathrm{t}$) resolution of the filter must set such that $(4 \cdot \Delta t \cdot D)/h^2 < 1$ (see footnote[5] and Thygesen et al., 2009).

**C. Likelihood.** Third, we update the prediction for the individual's location at the next time step with the likelihood $f_{ij}(\boldsymbol{y}) = f(\boldsymbol{y} \mid \boldsymbol{s}_{ij})$, thus providing an estimate of the partial marginal that accounts for the movement process and the data at time $t$:

$$P_{ij,t+1}(\boldsymbol{y}_{1:t+1}) = \frac{1}{Z(\boldsymbol{y}_{1:t+1})}\, f_{ij}(\boldsymbol{y}_{t+1})\, P_{ij,t+1}(\boldsymbol{y}_{1:t}), \qquad \text{eqn 14}$$

where $Z(\boldsymbol{y}_{1:t+1}) = \sum_{ij} P_{ij,t+1}(\boldsymbol{y}_{1:t})\, f_{ij}(\boldsymbol{y}_{t+1})$ is the normalisation constant. As the spatial ($h$) and temporal ($\Delta \mathrm{t}$) resolution tend to zero (with $\Delta t / h^2 \rightarrow 0$), we recover the continuous theory (eqn 10) from the discretised version presented above.

[5] `Wahoo.jl` users only need to ensure that the movement standard deviation ($\sigma_M$ or `movement_std` in the code) is expressed in units of observational time intervals. The filter steps along the timeline defined by the observations. We automatically compute $D$ and $\Delta t$ such that $(4 \cdot \Delta t \cdot D)/h^2 < 1$ and run the convolution as many times as necessary per time step. We refer to convolution steps within time steps as 'hops'.

The time complexity of the forward filter is $O(NT)$, where $N$ is the number of grid cells.

##### 1.2.2. Smoothing

Smoothing recursively re-weights probabilities from the filter to account for all data. That is, smoothing uses estimates of the partial marginal distributions, $f(\boldsymbol{s}_t \mid \boldsymbol{y}_{1:t})$, to approximate the full marginals, $f(\boldsymbol{s}_t \mid \boldsymbol{y}_{1:T})$. This is done recursively using the identity

$$\begin{aligned} f(\boldsymbol{s}_t \mid \boldsymbol{y}_{1:T}) &= f(\boldsymbol{s}_t \mid \boldsymbol{y}_{1:t}) \int f(\boldsymbol{s}_{t+1} \mid \boldsymbol{s}_t) \frac{f(\boldsymbol{y}_{t+1:T} \mid \boldsymbol{s}_{t+1})\, f(\boldsymbol{y}_{1:t})}{f(\boldsymbol{y}_{1:T})} d\boldsymbol{s}_{t+1} \\ &= f(\boldsymbol{s}_t \mid \boldsymbol{y}_{1:t}) \int f(\boldsymbol{s}_{t+1} \mid \boldsymbol{s}_t) \frac{f(\boldsymbol{s}_{t+1} \mid \boldsymbol{y}_{1:T})}{f(\boldsymbol{s}_{t+1} \mid \boldsymbol{y}_{1:t})} d\boldsymbol{s}_{t+1}. \end{aligned} \qquad \text{eqn 15}$$

The distribution $f(\boldsymbol{s}_t \mid \boldsymbol{y}_{1:T})$ is approximated on the grid recursively from $t = T$ to $t = 1$ via

$$P_{ij,t}(\boldsymbol{y}_{1:T}) = \begin{cases} P_{ij,t}(\boldsymbol{y}_{1:t}) & \text{if } t = T \\ P_{ij,t}(\boldsymbol{y}_{1:t}) \sum_{i'j'} P_{i'j' \mid ij} \dfrac{P_{i'j',t+1}(\boldsymbol{y}_{1:T})}{P_{i'j',t+1}(\boldsymbol{y}_{1:t})} & \text{otherwise,} \end{cases} \qquad \text{eqn 16}$$

where

$$P_{ij,t}(\boldsymbol{y}_{1:t}) \sum_{i'j'} P_{i'j' \mid ij} \frac{P_{i'j',t+1}(\boldsymbol{y}_{1:T})}{P_{i'j',t+1}(\boldsymbol{y}_{1:t})} = \sum_{r,s \,\in\, \{0,\pm 1\}} K_{rs} \frac{P_{i+r,j+s,t+1}(\boldsymbol{y}_{1:T})}{P_{i+r,j+s,t+1}(\boldsymbol{y}_{1:t})} \qquad \text{eqn 17}$$

and $K_{rs}$ is the ‘mirror image’ of $H_{rs}$, as $K_{rs} = P_{i+r,j+s,ij} = P_{ij,i-r,j-s} = H_{-r,-s}$, due to the translational invariance of the movement (diffusion) kernel. In practice, this approximation is achieved in two stages.

**A. Initialisation.** At $t = T$, set $P_{ij,t}(\boldsymbol{y}_{1:T}) = P_{ij,t}(\boldsymbol{y}_{1:t})$.

**B. Backward pass.** At each preceding time step, reweight the partial marginal $P_{ij,t}(\boldsymbol{y}_{1:t})$ from the filter by convolving the ratio of the full marginal distribution for the ‘previous’ (next) time step, $P_{i'j',t+1}(\boldsymbol{y}_{1:T})$ and the corresponding distribution from the forward filter at that time step, $P_{i'j',t+1}(\boldsymbol{y}_{1:t})$. The ratio $P_{i'j',t+1}(\boldsymbol{y}_{1:T})/P_{i'j',t+1}(\boldsymbol{y}_{1:t})$ effectively accounts for the future observation(s) at time $t$, which are then smoothed backwards in time in line with the reversed movement model.

For a diagrammatic explanation of the implementation of this algorithm, we direct the reader to the `Wahoo.jl` package documentation.

The overall occupancy distribution is defined by the normalised sum of all $P_{ij,t}(\boldsymbol{y}_{1:T})$ values across the grid:

$$P_{ij}(\boldsymbol{y}_{1:T}) = \frac{1}{T}\sum_{t=1}^{T} P_{ij,t}(\boldsymbol{y}_{1:T}). \qquad \text{eqn 18}$$

As in the filter, the time complexity of smoothing also scales linearly with the number of pixels $\big(O(NT)\big)$, but each time step is more expensive[6].

##### 1.2.3. Sampling trajectories

Trajectories are samples from the joint distribution $f(\boldsymbol{s}_{1:T} \mid \boldsymbol{y}_{1:T})$. Sampling trajectories is a recursive procedure that sweeps backwards in time. At each time step, we sample a position $\boldsymbol{s}_{t+1}$ and then condition on both that position and the data $\boldsymbol{y}_{1:t}$, according to the equation:

$$f(\boldsymbol{s}_t \mid \boldsymbol{y}_{1:t}, \boldsymbol{s}_{t+1}) = \frac{f(\boldsymbol{s}_t \mid \boldsymbol{y}_{1:t})\, f(\boldsymbol{s}_{t+1} \mid \boldsymbol{s}_t)}{f(\boldsymbol{s}_{t+1} \mid \boldsymbol{y}_{1:t})}. \qquad \text{eqn 19}$$

In discrete terms, we express the algorithm as follows.

A. **Initialisation.** At $t = T$, sample a cell, denoted $S_{ij}$, from $P_{ij,T}(\boldsymbol{y}_{1:t})$.

B. **Backward pass.** At each preceding time step, denote the cell sampled at time $t + 1$ as $S_{i'j'}$. Recursively compute the product of (i) the individual's possible locations at time $t$, given the movement capacity from $S_{i'j}$, obtained via convolution, and (ii) the data, according to the equation

$$w_{ij} \propto P_{ij,t}(\boldsymbol{y}_{1:t})\, P_{ij \mid i'j'}, \qquad \text{eqn 20}$$

where $w_{ij}$ denotes normalised weights[7]. Then sample a cell $S_{ij}$ at time $t$ via $w_{ij}$ and continue the recursion.

[6] In the worked example in this paper, we observed that filtering was nearly five times faster than smoothing on the GPU (1.5 times faster on the CPU).

[7] As for the filter, note that it may be necessary to repeat the convolution multiple times (hops) per time step. `Wahoo.jl` handles this as required.

The computational cost of the sampling one trajectory via convolution over the grid is similar to the cost of smoothing.

##### 1.2.4. Parameter estimation via the likelihood

The above inference algorithm estimates states, assuming static parameters are specified from prior analyses, domain knowledge and literature. In theory, it is possible to estimate static parameters over multiple filter runs via the likelihood $L(\boldsymbol{\theta})$ of the observations, given the parameters $\boldsymbol{\theta}$. The likelihood is derived from the joint distribution $f(\boldsymbol{y}_{1:T}, \boldsymbol{s}_{1:T} \mid \boldsymbol{\theta})$ by integrating out the latent states; i.e., $L(\boldsymbol{\theta}) = \int f(\boldsymbol{y}_{1:T}, \boldsymbol{s}_{1:T} \mid \boldsymbol{\theta})\, d\boldsymbol{s}_{1:T}$, where the joint distribution is given by the product

$$f(\boldsymbol{y}_{1:T}, \boldsymbol{s}_{1:T} \mid \boldsymbol{\theta}) = \prod_{t=1}^{T} f(\boldsymbol{y}_t \mid \boldsymbol{s}_t, \boldsymbol{\theta})\, f(\boldsymbol{s}_t \mid \boldsymbol{s}_{t-1}, \boldsymbol{\theta}) \qquad \text{eqn 21}$$

and $f(\boldsymbol{s}_1 \mid \boldsymbol{s}_0, \boldsymbol{\theta})$ is to be read as $f(\boldsymbol{s}_1)$. We compute the likelihood from the product of the normalisation denominators $Z(\boldsymbol{y}_{1:t})$ from the forward filter for $t = 1, \dots, T$:

$$L(\boldsymbol{\theta}) = \prod_{t=1}^{T} Z(\boldsymbol{y}_{1:t}). \qquad \text{eqn 22}$$

This can be shown by induction (applying Bayes' rule $T$ times):

$$\begin{aligned} Z(\boldsymbol{y}_{1:T}) &= \int f(\boldsymbol{y}_T \mid \boldsymbol{s}_T)\, f(\boldsymbol{s}_T \mid \boldsymbol{s}_{T-1})\, f(\boldsymbol{s}_{T-1} \mid \boldsymbol{y}_{1:T-1})\, d\boldsymbol{s}_T\, d\boldsymbol{s}_{T-1} \\ &= \frac{1}{Z(\boldsymbol{y}_{1:T-1})} \int d\boldsymbol{s}_T\, d\boldsymbol{s}_{T-1}\, d\boldsymbol{s}_{T-2}\, f(\boldsymbol{y}_T \mid \boldsymbol{s}_T)\, f(\boldsymbol{y}_{T-1} \mid \boldsymbol{s}_{T-1}) \\ &\qquad \times f(\boldsymbol{s}_T \mid \boldsymbol{s}_{T-1})\, f(\boldsymbol{s}_{T-1} \mid \boldsymbol{s}_{T-2})\, f(\boldsymbol{s}_{T-2} \mid \boldsymbol{y}_{1:T-2})\; d\boldsymbol{s}_T\, d\boldsymbol{s}_{T-1}\, d\boldsymbol{s}_{T-2} \\ &= (\dots) \\ &= L(\boldsymbol{\theta}) \prod_{t=1}^{T-1} \frac{1}{Z(\boldsymbol{y}_{1:t})}. \end{aligned} \qquad \text{eqn 23}$$

#### 2. Simulation analyses

##### 2.1. Example workflow

In the Main Text, we illustrate model-based inference with `Wahoo.jl` via simulation. Our illustration is motivated by research on flapper skate (*Dipturus intermedius*) in Scotland (Lavender et al., 2021b, 2021a, 2023; Lavender, Scheidegger, Albert, Biber, Aleynik, et al., 2025). The study area is defined by a 100 x 100 m bathymetry grid[8]. Within this area, we formerly deployed an array of 48 acoustic receivers and tagged skate with acoustic and archival depth tags to study their movements. Here, we leverage insights from previous work to simulate movements and acoustic and archival depth observations for a hypothetical individual[9]. These simulated data are then used to illustrate the `Wahoo.jl` workflow[10].

**Movements.** We simulated the movements of an individual within the study area at a time resolution of two minutes over a one-month period. The individual's initial location was sampled randomly within the area $A$ spanned by receivers:

$$\boldsymbol{s}_{t=1} \sim U(A). \qquad \text{eqn 24}$$

Subsequent movements were simulated from a Gaussian random walk model

$$\boldsymbol{s}_{t+1} \mid \boldsymbol{s}_t \sim N(\boldsymbol{s}_t, 100^2) \qquad \text{eqn 25}$$

with the condition that the individual cannot move onto land. The standard deviation was chosen in line with previous research (Lavender, Scheidegger, Albert, Biber, Aleynik, et al., 2025).

**Acoustic observations.** At each time step, we simulated acoustic observations (0, 1) by sampling from a Bernoulli distribution

$$y_{t,k}^{(A)} \,\Big|\, \boldsymbol{s}_t \sim \text{Bernoulli}\left(p_{t,k}(\boldsymbol{s}_t)\right) \qquad \text{eqn 26}$$

with a detection probability function

$$p_{t,k}(\boldsymbol{s}_t) = \left(1 + e^{-(4-0.0094\times|\boldsymbol{s}_t-\boldsymbol{r}_k|)}\right)^{-1} \qquad \text{eqn 27}$$

derived from range-testing data (Lavender, Scheidegger, Albert, Biber, Aleynik, et al., 2025).

[8] The bathymetric grid is based on a 5 x 5 m data product provided by Howe et al. (2014).
[9] Movements and observations were simulated using the `patter` package (Lavender, Scheidegger, Albert, Biber, Illian, et al., 2025b).
[10] By basing our example on simulated data, we can compare simulated and inferred patterns and explore the behaviour of the grid-based inference procedure.

**Depth observations.** Depths were sampled from a truncated Gaussian distribution, centred on the seabed

$$y_t^{(D)} \mid \boldsymbol{s}_t \sim \text{Truncated}(N(b(\boldsymbol{s}_t)), 12.7527), 0, 350) \qquad \text{eqn 28}$$

with a standard deviation that encapsulates bathymetric accuracy, tidal ranges, storm surges and tag accuracy[11] (Lavender, Scheidegger, Albert, Biber, Aleynik, et al., 2025).

**Inference.** Workflows written in `Julia` and `R` for the analysis of these data via `Wahoo.jl` are provided as supporting documents. We initialised the filter with a uniform distribution over the sea and used the correct data-generating movement and observation models to perform inference for the latent locations[12]. This provides a 'best-case' benchmark for the performance of the grid-based filter in this study system. However, we also illustrate a sensitivity analysis that evaluates the consequences of parameter misspecification for inference of the latent locations (see §2.3). Routines were run on the GPU and the CPU to compare computation time.

#### 2.2. Validation analyses

We conducted a more extensive analysis to validate `Wahoo.jl` and illustrate how the grid-based filtering approach compares to a particle-based approach (implemented by `patter`)[13]. For this analysis, we simulated movements and observations for five individuals[14], following the workflow described in §2.1. We then analysed simulated observations using `Wahoo.jl` and

[11] The Gaussian model is based on the knowledge that skate are predominantly benthic animals (Lavender, Scheidegger, Albert, Biber, Aleynik, et al., 2025). The standard deviation accounts for uncertainty in the bathymetric depth, tidal elevation, storm surges and waves and archival tag accuracy. For our study system, these uncertainties were quantified by Lavender, Scheidegger, Albert, Biber, Aleynik, et al. (2025). We consider uncertainties of ± 10/2 m (bathymetry), ± 3/2 m (tidal elevation), ± 4.77/2 m (archival tag accuracy). We divide each source of uncertainty by two based on the assumption that 95% % of data are within ± 2 standard deviations of the mean. We account for the additional uncertainty introduced by aggregating bathymetric grid cells from 5 x 5 m to 100 x 100 m by computing the mean standard deviation in bathymetric depths within each 100 x 100 m grid cell (over all grid cells). This was 2.867705 m. Collectively, these sources of uncertainty suggest a parameterisation of $\sigma_D = 12.75270$ m. The tails of the normal distribution permit larger movements away from the seabed (allowing potential pelagic behaviour) with lower probability.

[12] In the simulation, we programmed that criteria that the fish cannot move onto land into the movement model. For model inference, we use an untruncated Gaussian random walk, enabling the movement process to be represented via convolution. During inference, the depth observation model $f\left(y_t^{(D)} \mid \boldsymbol{s}_t\right)$ ensures that fish cannot move onto land.

[13] The ruggedness of the bathymetric landscape and the precision of the depth observation model make this example a hard inference problem for sampling methods.

[14] Following Lavender, Scheidegger, Albert, Biber, Aleynik, et al. (2025), we repeated the simulation iteratively until we generated a movement path associated with sufficient observations to merit modelling. (Two attempts were necessary for two individuals.)

`patter`. Inference with `Wahoo.jl` was performed as in §2.1. For inference with `patter`, we conducted a preliminary analysis with 20,000 particles for the filter and 1,000 for the smoother to gauge computation time. The full analysis used 100,000 particles for the filter and 2,000 particles for smoothing. Movement and observation models were specified as in `Wahoo.jl`. Resampling was implemented when the effective sample size was ≤ 1000 particles. The implementation was performed in single-threaded model with batching. Following inference, we visually compared simulated and inferred occupancy distributions from `Wahoo.jl` and `patter`. We also analysed `patter` diagnostics to investigate sampling efficacy.

##### 2.3. Sensitivity analyses

In the above analyses (§2.1–2), we performed inference for the latent locations using the correct data-generating models. This is a best-case scenario. In practice, there is typically a degree of uncertainty in static parameters. We therefore illustrate a sensitivity analysis, re-analysing the datasets simulated above with mis-specified algorithm parameterisations[15]. In each re-analysis, we performed inference with one mis-specified movement/observation model while holding other models at the correct (data-generating) settings. Following Lavender, Scheidegger, Albert, Biber, Aleynik, et al., (2025), for each model we considered restrictive and flexible parameterisations (Table S2). As in the validation analysis, we then visually compared occupancy distributions for correct and mis-specified algorithm applications. As a quantitative metric, we computed the Jensen-Shannon divergence between the best occupancy distributions and each occupancy distribution derived from the mis-specified algorithm run.

[15] As noted in §1.2.4 and elsewhere, in theory it is possible to latent locations and static parameters if the computations can be managed. However, in acoustic telemetry systems, data are typically limited and we prefer to focus inference on the latent locations under plausible settings for the movement and observation models informed by domain knowledge.
