## Supporting Figures for "Animal geolocation with convolution algorithms in Julia and R via Wahoo.jl"

Fig. S1. **Validation analyses of `Wahoo.jl` with `patter` (overleaf).** Rows distinguish individuals. For each individual, the true occupancy distribution and the distributions reconstructed by `Wahoo.jl` (Wah) and `patter` (Pat) are shown. For `patter`, occupancy distributions were derived from two-filter smoothing (which uses a forward filter run and a backward filter run). The proportion of time steps for which proper smoothing and an effective sample size exceeding 500 was achieved is marked. (In the particle algorithms, if the two filters diverge, proper smoothing is not possible and `patter` simply retains 50 % of particles from each of the two filter runs.) Note the difference between `Wahoo.jl` and `patter` outputs associated with poor sampling efficacy in the final row[1].

[1] The particle filter was implemented with 100,000 particles; the smoother was implemented with 2,000 particles. During manuscript development, we observed the occupancy distributions remained broadly similar with fewer particles (20,000 in the filter and 1,000 in the smoother) though effective sample sizes were smaller.

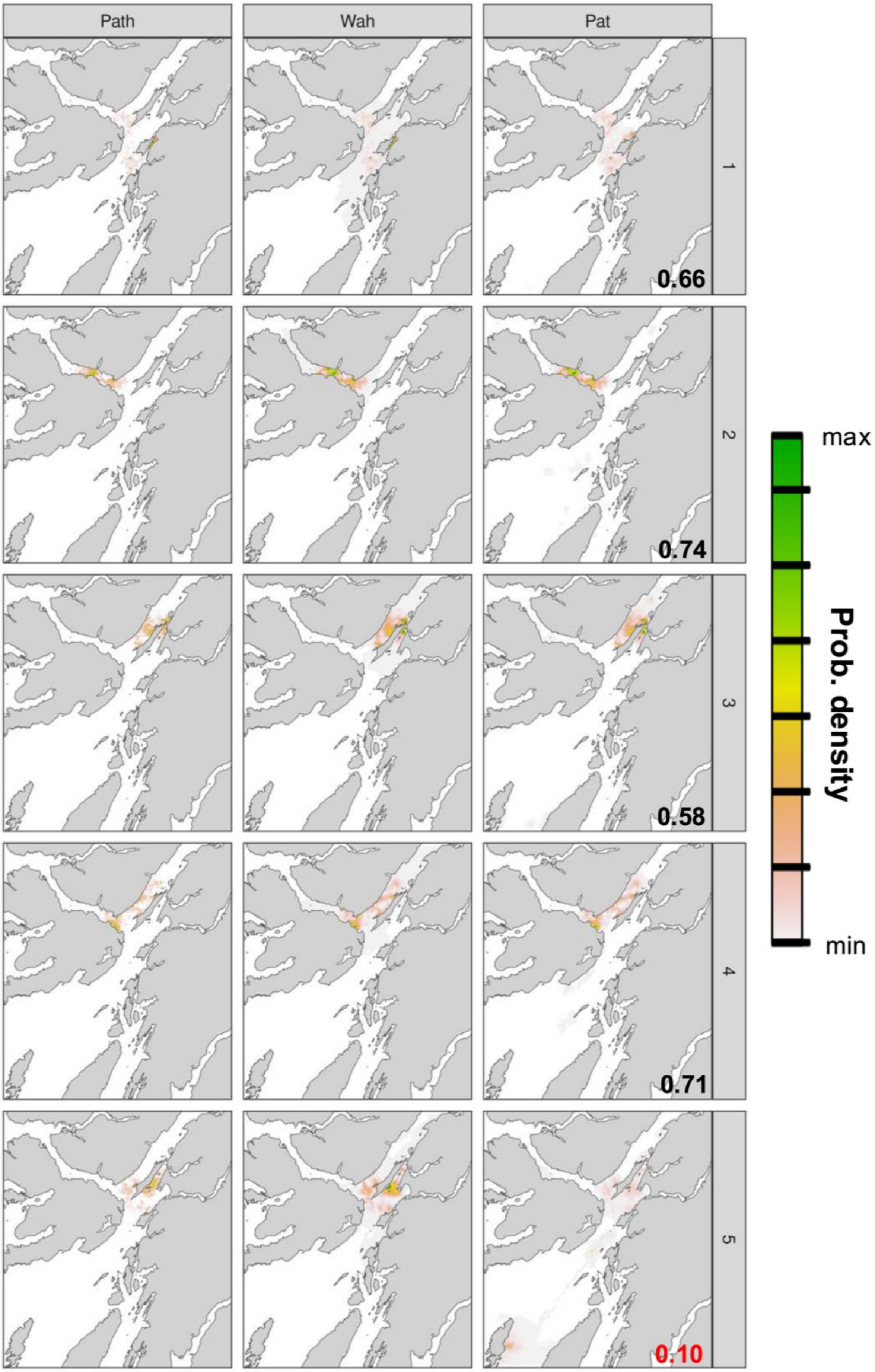
Path
Wah
Pat
1
2
3
4
5
0.66
0.74
0.58
0.71
0.10
max
Prob. density
min

**Fig. S2. Sensitivity analyses of `Wahoo.jl` and `patter` (overleaf).** Rows distinguish individuals. For each individual, the 'best' occupancy distribution is the distribution reconstructed using data-generating parameters (see Fig. S1). Occupancy distributions reconstructed by algorithm runs with mis-specified movement (M), acoustic (A) and depth (D) observation models are also shown. For each mis-specified model, we considered a restrictive (-) and flexible (+) parameterisation (the parameters of other models were held constant at the true values). Distributions reconstructed by `Wahoo.jl` (Wah) and `patter` (Pat) are shown. For a visualisation of model parameterisations, see Fig. 1. For parameter values, see Table S2.

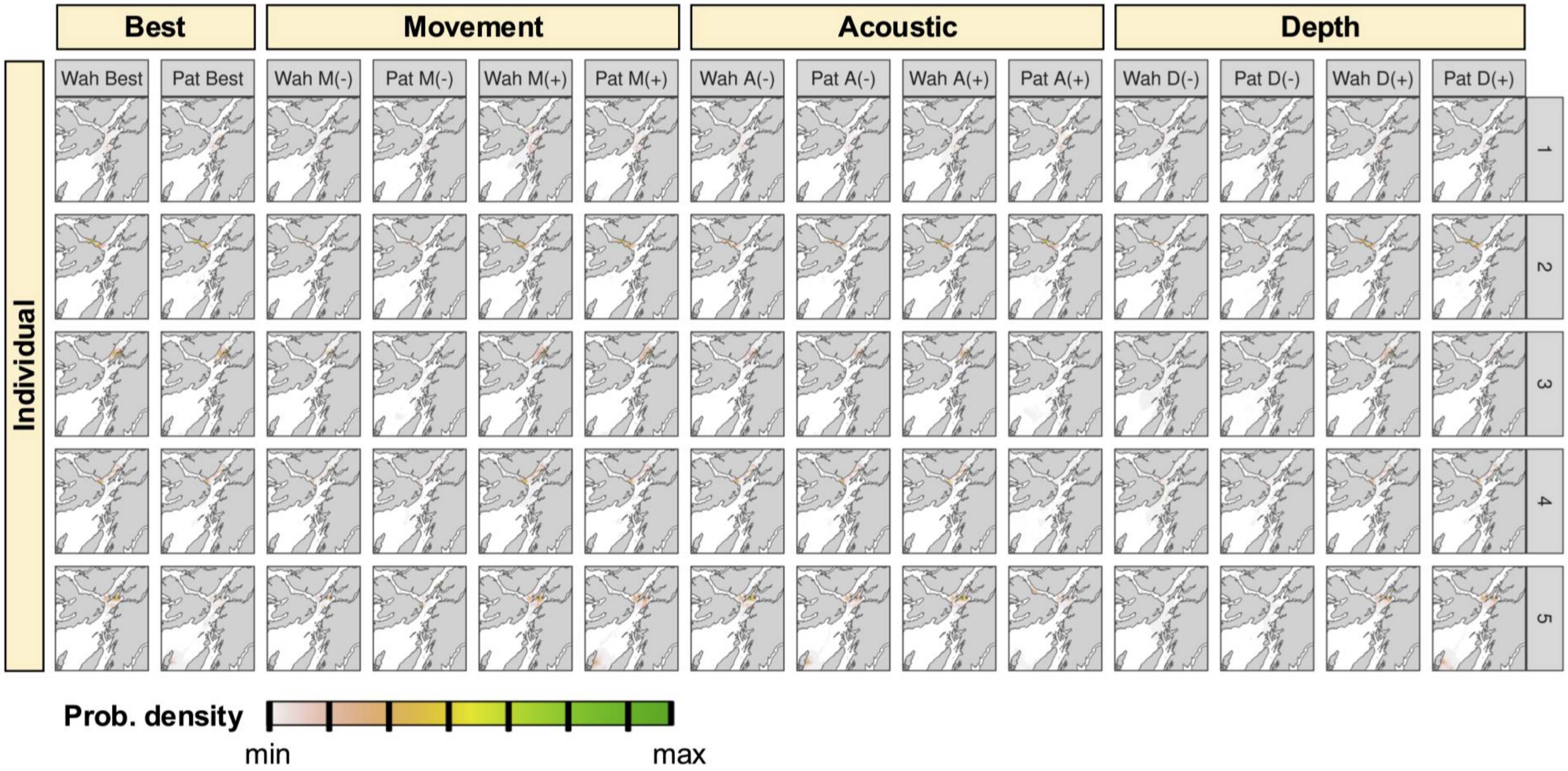
Best
Movement
Acoustic
Depth
Wah Best
Pat Best
Wah M(-)
Pat M(-)
Wah M(+)
Pat M(+)
Wah A(-)
Pat A(-)
Wah A(+)
Pat A(+)
Wah D(-)
Pat D(-)
Wah D(+)
Pat D(+)
Individual
1
2
3
4
5
Prob. density
min
max

**Fig S3. Sensitivity scores for `Wahoo.jl` and `patter`.** Panels distinguish individuals. Each panel shows, for both `Wahoo.jl` and `patter`, Jensen–Shannon divergences between the best occupancy distribution for that package and the corresponding distributions reconstructed by mis-specified algorithm runs[2]. Generally higher divergences for `patter` indicate additional numerical instability associated with parameter misspecification compared to `Wahoo.jl`.

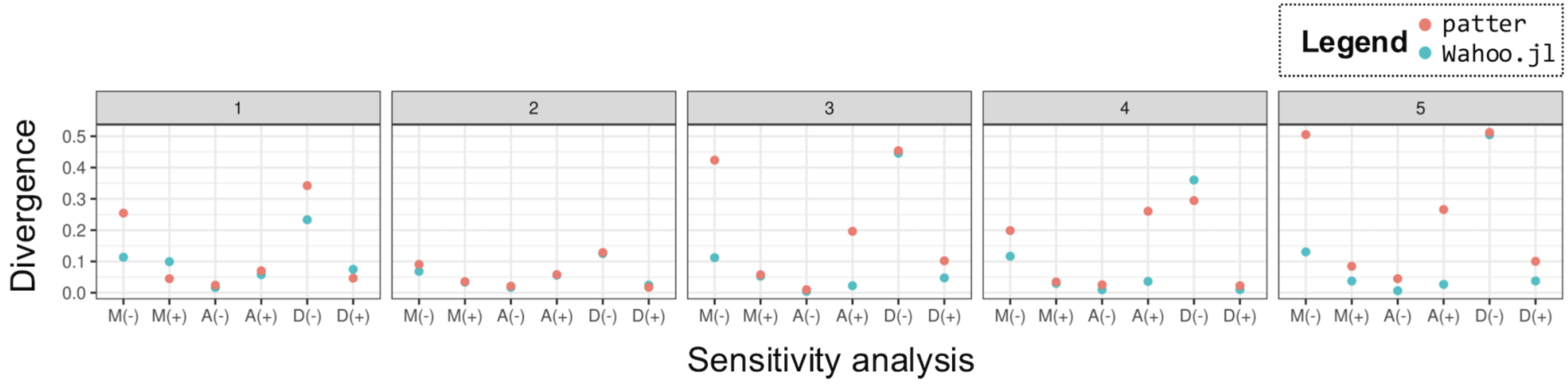


[2] As described in Fig. S2, in sensitivity analyses, we reimplemented `Wahoo.jl` and `patter` routines using mis-specified movement (M), acoustic (A) and depth (D) observation models. For each mis-specified model, we considered a restrictive (-) and flexible (+) parameterisation (the parameters of other models were held constant at the true values).
