## Supporting Tables for "Animal geolocation with convolution algorithms in Julia and R via Wahoo.jl"

**Table S1. A mapping of the notation from this manuscript to Thygesen et al. (2009), who outlined the grid-based filtering methodology.**

| Distribution | Notation | | |
|---|---|---|---|
| | **Generic** | **Discrete** | **Thygesen et al. (2009)** |
| **'Prior'** | $f(\boldsymbol{s}_{t+1} \mid \boldsymbol{y}_{1:t}) = f(\boldsymbol{s}_{t+1} \mid \boldsymbol{s}_t)\, f(\boldsymbol{s}_t \mid \boldsymbol{y}_{1:t})$ | $P_{ij,t+1}(\boldsymbol{y}_{1:t}) = \sum_{i'j'} P_{ij \mid i'j'}\, P_{i'j',t}(\boldsymbol{y}_{1:t})$ | $\Phi(t_{k+1} \mid t_k) = H(t_k, t_{k+1}) * \Phi(t_k \mid t_k)$ |
| → Movement model/transition probabilities | $f(\boldsymbol{s}_{t+1} \mid \boldsymbol{s}_t)$ | $P_{ij \mid i'j'}$ or $H_{rs}$ | $H(t_k, t_{k+1})$ |
| → Likelihood | $f(\boldsymbol{y}_t \mid \boldsymbol{s}_t)$ | $f_{ij}(\boldsymbol{y}_t)$ | $\Phi(t_k \mid t_k)$ |
| **Partial marginal**, estimated by the forward filter | $f(\boldsymbol{s}_{t+1} \mid \boldsymbol{y}_{1:t+1}) = \frac{1}{Z(\boldsymbol{y}_{1:t+1})} f(\boldsymbol{y}_{t+1} \mid \boldsymbol{s}_{t+1})\, f(\boldsymbol{s}_{t+1} \mid \boldsymbol{y}_{1:t})$ | $P_{ij,t+1}(\boldsymbol{y}_{1:t+1}) = \frac{1}{Z(\boldsymbol{y}_{1:t+1})} f_{ij}(\boldsymbol{y}_{t+1})\, P_{ij,t+1}(\boldsymbol{y}_{1:t})$ | $\Phi(t_{k+1} \mid t_{k+1}) \propto H(t_k, t_{k+1}) * \Phi(t_k \mid t_k) \times L(t_{k+1})$ |
| **Full marginal,** estimated by smoothing | $f(\boldsymbol{s}_t \mid \boldsymbol{y}_{1:T}) = f(\boldsymbol{s}_t \mid \boldsymbol{y}_{1:t}) \int f(\boldsymbol{s}_{t+1} \mid \boldsymbol{s}_t) \frac{f(\boldsymbol{s}_{t+1} \mid \boldsymbol{y}_{1:T})}{f(\boldsymbol{s}_{t+1} \mid \boldsymbol{y}_{1:t})} d\boldsymbol{s}_{t+1}$ | $P_{ij,t}(\boldsymbol{y}_{1:T}) = \begin{cases} P_{ij,t}(\boldsymbol{y}_{1:t}) & \text{if } t = T \\ P_{ij,t}(\boldsymbol{y}_{1:t}) \sum_{i'j'} P_{i'j' \mid ij} \frac{P_{i'j',t+1}(\boldsymbol{y}_{1:T})}{P_{i'j',t+1}(\boldsymbol{y}_{1:t})} & \text{otherwise} \end{cases}$ | $\Phi(t_{k+1} \mid \infty) = \begin{cases} \Phi(t_k \mid t_k) & \text{if } k = N \\ \Phi(t_k \mid t_k) \times \left[K(t_k, t_{k+1}) * \frac{\Phi(t_{k+1} \mid \infty)}{\Phi(t_{k+1} \mid t_k)}\right] & \text{otherwise} \end{cases}$ |

**Table S2. A summary of the parameter values used in analyses.** The example and validation analyses were run with the true (data-generating) parameters. We motivated these analyses by choosing parameters previously derived for an example species: the flapper skate (*Dipturus intermedius*). In sensitivity analyses, we re-analysed simulated datasets with mis-specified movement (M), acoustic (A) and depth (D) observation models. For each mis-specified model, we considered a restrictive (-) and flexible (+) parameterisation (the parameters of other models were held constant at the true values). For a visualisation of all model parameterisations, see Fig. 1.

| **Parameter ID** | **Sensitivity** | $\sigma_M$, `movement_std` | $\alpha$, `receiver_alpha` | $\beta$, `receiver_beta` | $\sigma_D$, `depth_std` |
|---|---|---|---|---|---|
| 1 | True | 100.0 | 4.0 | -0.00940 | 12.75270 |
| 2 | M(-) | 50.0 | 4.0 | -0.00940 | 12.75270 |
| 3 | M(+) | 150.0 | 4.0 | -0.00940 | 12.75270 |
| 4 | A(-) | 100.0 | 3.0 | -0.01175 | 12.75270 |
| 5 | A(+) | 100.0 | 5.0 | -0.00705 | 12.75270 |
| 6 | D(-) | 100.0 | 4.0 | -0.00940 | 6.376350 |
| 7 | D(+) | 100.0 | 4.0 | -0.00940 | 19.12905 |
