## Supporting Code (R) for "Animal geolocation with convolution algorithms in Julia and R via Wahoo.jl"

### An example workflow for `Wahoo.jl` in `R`

#### Introduction

This is an example `R`–`Wahoo.jl` workflow motivated by our research on flapper skate (*Dipturus intermedius*) in Scotland. We simulated an example movement trajectory for a hypothetical individual and acoustic and archival depth observations arising from the simulated path (using the `patter` package). The workflow below reconstructs simulated movements from `R` via the `Julia` package `Wahoo.jl`.

#### Set up

We begin with boilerplate set up: we load selected `R` packages; start a `Julia` session; activate a local `Julia` environment; and load necessary `Julia` packages. We use the `JuliaSwitch` package as our `R`–`Julia` interface. This package provides a common syntax for interacting with `Julia` from `R` via `JuliaCall` or `JuliaConnectoR`. `JuliaCall` uses a `C` interface to connect `R` and `Julia`, while `JuliaConnectoR` uses Transfer Control Protocol, which can be slower but is more stable across different system configurations (especially when dynamic libraries are involved). Using `JuliaSwitch`, we can run `Julia` code via a `julia_*()` helper function or `julia_cmd_line()`/ `julia_cmd_block()`; push objects to `Julia` using `julia_push()`; and pull them back via `julia_pull()`. For a richer functional experience, see the `JuliaConnectoR` documentation.

```
# Start time
t1 <- Sys.time()

# Load R packages
library(data.table)
library(dtplyr)
library(dplyr, warn.conflicts = FALSE)
library(here)
library(glue)
library(JuliaSwitch)
library(patter)
library(rhdf5)

# Set options
options(patter.verbose = FALSE, terra.pal = rev(terrain.colors(256L)))
```

```
# Start Julia
julia_backend("JuliaCall")
julia_start()
#>    Julia    version    1.11.6    at    location    /Users/lavended/.julia/juliaup/
```

```
julia-1.11.6+0.aarch64.apple.darwin14/bin will be used.
#> Loading setup script for JuliaCall...
#> Finish loading setup script for JuliaCall.
```

```
# Activate local Julia environment
julia_pkg_activate(here())

# Load Julia packages
julia_using("CSV")
julia_using("DataFrames")
julia_using("Dates")
julia_using("GeoArrays")
julia_using("HDF5")
julia_using("JLD2")
julia_using("NNlib")
julia_using("Plots")
julia_using("SpecialFunctions")

# Load Wahoo
julia_using("Wahoo")
```

```
# (optional) Import CUDA and cuDNN
# * If imported, Wahoo.jl will exploit GPU acceleration
# * Currently, Wahoo.jl only supports CUDA-compatible GPUs
julia_import("CUDA")
julia_import("cuDNN")
```

### Study system

#### Datasets

The next step is to load relevant datasets. We will load datasets into `R` and then push them to `Julia`. We load:

- **Study area bathymetric raster**. We use a 5 x 5 m raster produced by Lavender et al. (2025), downscaled to 100 x 100 m. Depth is coded in metres. Land is coded as −1 m. The coordinate reference system is UTM 29N.
- **Study timeline**. We consider a one-month timeline with a resolution of 2-minutes. This is the time resolution at which movements and acoustic and archival depth observations were simulated.
- **Simulated path**. This is the simulated trajectory, with one position per time step.
- **Acoustic receiver positions.** We use receiver positions for the receiver array deployed by the Movement Ecology of Flapper Skate (MEFS) project.
- **Acoustic observations.** These are defined in a `data.frame`. Each row is a time step. Each column is a receiver. Cells define detections (1), non-detections (0) or inactive receivers (NA).

- **Depth observations.** These are defined in a `data.frame`. Each row is a time step. Numbers define individual depths (m).

```
# Define study area
bathy <- terra::rast(here("data/input/map-wahoo-100m.tif"))
```

```
# Define timeline
timeline <- seq(as.POSIXct("2025-01-01 00:00:00", tz = "UTC"),
                as.POSIXct("2025-01-31 23:58:00", tz = "UTC"),
                by = "2 mins")
head(timeline, 3)
#> [1] "2025-01-01 00:00:00 UTC" "2025-01-01 00:02:00 UTC"
#> [3] "2025-01-01 00:04:00 UTC"
```

```
# Define simulated path
path <-
  read.csv(here("data/input/sim/example/path.csv"))
head(path, 3)
#>   individual_id           timestamp timestep map_value        x       y
#> 1             1          2025-01-01        1  50.17444 708949.6 6262984
#> 2             1 2025-01-01 00:02:00        2  59.46473 708803.3 6262821
#> 3             1 2025-01-01 00:04:00        3  42.13493 708781.5 6262870
```

```
# Define receiver positions
moorings_df <-
  read.csv(here("data/input/sim/example/moorings-wahoo.csv"))
head(moorings_df, 3)
#>   receiver_id receiver_x receiver_y
#> 1           2   719404.5    6273496
#> 2           3   706472.2    6254001
#> 3           4   709757.8    6267727
```

```
# Define acoustic observations (0, 1)
acoustic_obs_df <-
  read.csv(here("data/input/sim/example/acoustics-wahoo.csv"))
acoustic_obs_df[1:3, 1:10]
#>   X2 X3 X4 X5 X7 X8 X9 X10 X11 X12
#> 1  0  0  0  0  0  0  0   0   0   0
#> 2  0  0  0  0  0  0  0   0   0   0
#> 3  0  0  0  0  0  0  0   0   0   0
```

```
# Define depth observations
depth_obs_df <-
  read.csv(here("data/input/sim/example/depths-wahoo.csv"))
```

```r
head(depth_obs_df, 3)
#>      depth
#> 1 56.15854
#> 2 59.24517
#> 3 43.97440
```

```r
# Push data to Julia
julia_push("bathy", bathy)
julia_push("timeline", timeline)
julia_push("moorings_df", moorings_df)
julia_push("acoustic_obs_df", acoustic_obs_df)
julia_push("depth_obs_df", depth_obs_df)
```

#### Map

This is a map of the study area with the simulated path and receiver array added:

```r
terra::plot(bathy)
patter:::add_sp_path(path$x, path$y,
                     length = 0, lwd = 0.5)
points(moorings_df$receiver_x, moorings_df$receiver_y,
       col = "red", lwd = 1.25)
```

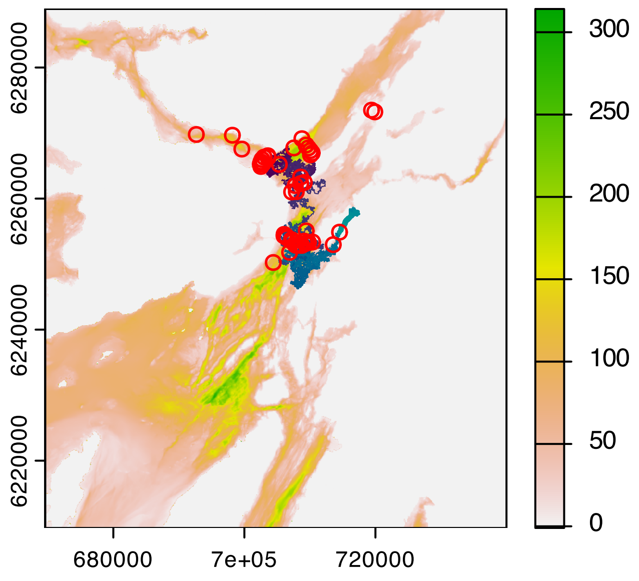

#### Model formulation

Now we formulate our state-space model. This comprises two sub-models: a movement model and a stack of observation models (one for each sensor).

##### Movement model

`Wahoo.jl` implements a Gaussian random walk (diffusion) movement model via two-dimensional convolution. We set the standard deviation of the movement model as `movement_std`. We select `movement_std = 100` m based on previous research (Lavender et al., 2025). For this workflow, we treat `movement_std` (and the static parameters in the observation models) as known and focus inference on the latent states. (However, we note that in principle it is possible to estimate static parameters via the log-likelihood if data are sufficient and the computations can be managed.)

```
julia_cmd_line('movement_std = 100.0')
```

##### Observation models

Observation models provide the link between the latent states and our observations. `Wahoo.jl` expects observations to be provided in a `Vector`, with one element for each sensor (e.g., receiver). Each element is a time series of observations for that sensor (with one observation per time step). For each sensor, a corresponding observation model should be provided. `Wahoo.jl` includes built-in observation models or we can define custom models with the signature:

```
function p_obs_*(signals, t::Int, depth::Number, distance::Number; kwargs...)
    _p_obs_*(signals[t], depth, distance; kwargs...)
end
```

###### Acoustic observation model

The acoustic observation model acts on acoustic observations. We provide acoustic observations in a `Vector`, with one element for each sensor. Each element is a time series of acoustic observations (0, 1 or −1) for that sensor. We use a Bernoulli observation model in which the probability of a detection is assumed to decline logistically with distance from a receiver. We use a model in which the rate of decline is controlled by two parameters, `receiver_alpha` and `receiver_beta`, informed by prior research (Lavender et al., 2025).

```
# Define a Vector with receiver positions:
julia_cmd_line('
  acoustic_pos = tuple.(moorings_df.receiver_x, moorings_df.receiver_y)
')
```

```
# Collate acoustic observations in a Vector
# * Code observations as:
#   - 1 (detection)
#   - 0 (non-detection)
```

```
#   - -1 (no signal: inactive receiver)
# * We define a Vector of Vectors (each element is a sensor)
julia_cmd_block(
'
acoustic_obs = coalesce.(transpose(Array(acoustic_obs_df)), -1)
acoustic_obs = [acoustic_obs[i, :] for i in 1:size(acoustic_obs, 1)]
'
)
```

```
# Define acoustic observation models
julia_cmd_block(
'
# Define acoustic observation model function
function p_obs_acoustic(signals, t::Int, waterdepth::Number, distance::Number;
                        receiver_alpha = 4.0, receiver_beta = -0.00940)
    signal = signals[t]
    if signal == 0
        return 1 - NNlib.sigmoid(receiver_alpha + receiver_beta * distance)
    end
    if signal == 1
        return NNlib.sigmoid(receiver_alpha + receiver_beta * distance)
    end
    return one(waterdepth)
end

# Define a Vector of observation models (one for each sensor)
p_obs_acoustic_vect = [p_obs_acoustic for i in 1:length(acoustic_obs)];
'
)
```

##### Depth observation model

As we have depth observations, we also define a depth observation model. Depth observations are collated in a `Vector`, as for acoustic observations. We define a truncated Gaussian observation model, with a mean on the seabed (in line with the benthic habit of skate) and a standard deviation that captures uncertainty in the bathymetric and depth measurements (based on prior research).

```
# Collect depth observations in a Vector
julia_cmd_line('depth_obs = Vector(depth_obs_df.depth)')
```

```
# Define depth observation model
julia_cmd_block(
'
function p_obs_depth(signals, t::Int, waterdepth::Number, distance::Number;
                     depth_std = 12.75270)
    signal = signals[t]
```

```
    if waterdepth < 0
        return zero(waterdepth)
    else
        mu = waterdepth
        f_obs = (1.0 / (sqrt(2.0 * pi) * depth_std)) *
                   exp(-((signal - mu)^2) / (2.0 * depth_std^2))
        lwr = (0.0 - mu) / (sqrt(2.0) * depth_std)
        upr = (350.0 - mu) / (sqrt(2.0) * depth_std)
        Z = 0.5 * (1.0 + erf(upr)) - 0.5 * (1.0 + erf(lwr))
        return f_obs / Z
    end
end
'
)
```

#### Inference

Now we can run the inference routine. A single function, `track()`, is exported for this purpose. We provide `track()` with a probability distribution for the animal's initial location (coded as a `Matrix`). For simplicity, we select a uniform distribution over the marine area. We also supply our bathymetric raster and its spatial resolution, the time points at which we want to record probability distributions and our movement and observation models.

```
# Define probability distribution for the individual's initial location
p0 <- terra::deepcopy(bathy)
p0[p0 >= 0] <- 1
p0[p0 == -1] <- 0
julia_push("p0", p0)
julia_cmd_line("p0 = Matrix(p0)")
```

```
# Define save time points as an _integer_
# julia_cmd_line('tsave = 1:60:length(timeline)')
tsave <- seq(1L, length(timeline), by = 60L)
julia_push("tsave", tsave)
```

```
# Run inference
outfile <- here("data/output/sim/example/R/wahoo.h5")
if (!file.exists(outfile)) {

  # Run track()
  print("Running Wahoo.jl")
  julia_cmd_block(
    '
    res = track(pos_init           = p0,
                bathymetry         = bathy,
```

```
                 spatial_resolution = 100,
                 tsave              = tsave,
                 movement_std       = movement_std,
                 observations       = [depth_obs, acoustic_obs...],
                 observation_models = [p_obs_depth, p_obs_acoustic_vect...],
                 sensor_positions   = [nothing, acoustic_pos...],
                 filter             = true,
                 smoother           = true,
                 n_trajectories     = 1,
                 show_progressbar   = true,
                 precision          = Float64)
     ')

   # Examine summary
   # * track() returns a NamedTuple:
   # * res.pos_filter
   #   > Array{Float, 4}: Ny × Nx × 1 × time of Prob(s_t | y_{1...t})
   # * res.pos_smoother
   #   > Array{Float, 4}: Ny × Nx × 1 × time of Prob(s_t | y_{1...T})
   # * res.residence_dist
   #   > Matrix{Float}: Ny × Nx of 1/T Sum Prob(s_t | y_{1...T})
   # * res.trajectories
   #   > Vector of trajectories from Prob(s_{1...T} | y_{1...T})
   # * res.log_p
   #   > Vector of Prob(y_t)
   # * res.tsave
   #   > Vector of time points

   # (optional) Write output to file via HDF5
   julia_cmd_block(glue(
   '
   h5open("{outfile}", "w") do f
     f["timesteps"] = collect(res.tsave)
     f["pos_smoother"] = dropdims(res.pos_smoother, dims = 3)
     f["residence_dist"] = res.residence_dist
     isdefined(res, :pos_filter) &&
       (f["pos_filter"] = dropdims(res.pos_filter, dims = 3))
     f["log_p"] = res.log_p
     !isnothing(res.trajectories) &&
       foreach(enumerate(res.trajectories)) do (i, tr)
         f["trajectories/traj$(i)"] = tr
       end
   end
   '
   ))

 }
```

#### Analysis

We can analyse `Wahoo.jl`'s outputs using all of the standard tools available in the `R` ecosystem. As a starting point, below we illustrate how to read and plot the estimated residency distribution and sampled trajectories, which we compare against the simulated path. As an exercise, we invite you to deepen this investigation of algorithm outputs and diagnostics.

```r
# Read selected Wahoo outputs
wahoo_occ  <- h5read(outfile, "residence_dist")
wahoo_traj <- h5read(outfile, "trajectories")

# Compute occupancy for simulated path
bathy     <- terra::classify(bathy, cbind(-1, NA))
path_occ  <- map_pou(.map = bathy, .coord = path, .plot = FALSE)$ud

# Extract estimated occupancy
wahoo_occ <- terra::rast(wahoo_occ)
wahoo_occ <- terra::t(wahoo_occ)
terra::ext(wahoo_occ) <- terra::ext(bathy)
terra::crs(wahoo_occ) <- terra::crs(bathy)
wahoo_occ <- terra::mask(wahoo_occ, bathy)

# Extract trajectories
wahoo_traj <- lapply(seq_along(wahoo_traj), function(i) {
  # Extract array indices (ai)
  ai <- wahoo_traj[[i]]
  ay <- ai[1, ] # array row (y)
  ax <- ai[2, ] # array col (x)
  # Swap to raster indices
  rr <- ax
  rc <- ay
  # Define xy coordinates
  cells <- terra::cellFromRowCol(bathy, row = rr, col = rc)
  xy    <- terra::xyFromCell(bathy, cells)
  stopifnot(!any(is.na(terra::extract(bathy, xy)[, 1])))
  # Collate data.table
  data.table(path_id = i,
             timestep = seq_len(ncol(ai)),
             x        = xy[, 1],
             y        = xy[, 2])
}) |> rbindlist()

# Extract the first sampled trajectory
wahoo_traj_1 <- wahoo_traj[path_id == 1L, ]

# Visualise maps
pp <- par(mfrow = c(2, 2))
terra::plot(path_occ, main = "A, Simulated occupancy")
```

```
terra::plot(wahoo_occ, main = "B, Wahoo occupancy")
terra::plot(path_occ, main = "C, Simulated path")
patter:::add_sp_path(path$x, path$y, length = 0, lwd = 0.2)
terra::plot(wahoo_occ, main = "D, Wahoo path")
patter:::add_sp_path(wahoo_traj_1$x, wahoo_traj_1$y, length = 0, lwd = 0.2)
```

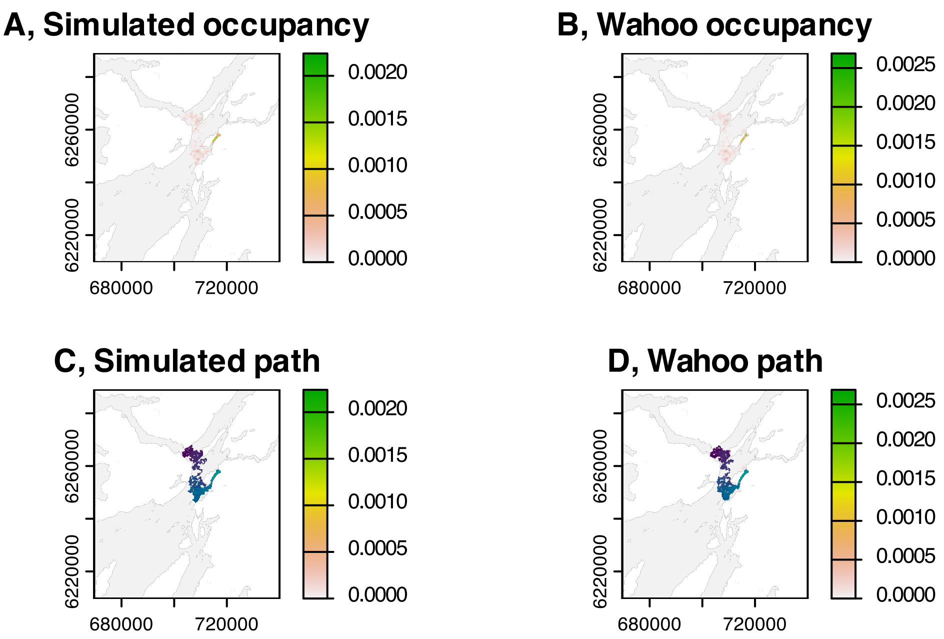


```
par(pp)
```

#### Close Julia

```
# Close Julia session
julia_stop()
#> JuliaCall does not support termination of the Julia session.
```

```
# Record render duration (mins)
t2 <- Sys.time()
difftime(t2, t1, units = "mins")
#> Time difference of 0.3206845 mins
```
