## Supporting Code (Julia) for "Animal geolocation with convolution algorithms in Julia and R via Wahoo.jl"

### An example workflow for `Wahoo.jl` in `Julia`

#### Introduction

This is an example `Wahoo.jl` workflow motivated by our research on flapper skate (*Dipturus intermedius*) in Scotland. We simulated an example movement trajectory for a hypothetical individual and acoustic and archival depth observations arising from the simulated path (using the `patter` package). The workflow below reconstructs simulated movements via the `Julia` package `Wahoo.jl`.

#### Set up

The first step is to activate our local `Julia` environment and load packages.

```
# Activate local Julia environment
import Pkg
Pkg.activate("..")

# Load Julia packages
using CSV
using DataFrames
using Dates
using GeoArrays
using JLD2
using NNlib
using Plots
using SpecialFunctions

# Load Wahoo
using Wahoo
```

```
  Activating project at `~/Documents/work/projects/move-fokker-plank/projects/
wahoo-flapper`
```

```
# (optional) Import CUDA and cuDNN
# * If imported, Wahoo.jl will exploit GPU acceleration
# * Currently, Wahoo.jl only supports CUDA-compatible GPUs
import CUDA
import cuDNN
```

#### Study system

#### Datasets

The next step is to load relevant datasets:

- **Study area bathymetric raster**. We use a 5 x 5 m raster produced by Lavender et al. (2025), downscaled to 100 x 100 m. Depth is coded in metres. Land is coded as −1 m. The coordinate reference system is UTM 29N.
- **Study timeline**. We consider a one-month timeline with a resolution of 2-minutes. This is the time resolution at which movements and acoustic and archival depth observations were simulated.
- **Simulated path**. This is the simulated trajectory, with one position per time step.
- **Acoustic receiver positions.** We use receiver positions for the receiver array deployed by the Movement Ecology of Flapper Skate (MEFS) project.
- **Acoustic observations.** These are defined in a `DataFrame`. Each row is a time step. Each column is a receiver. Cells define detections (1), non-detections (0) or inactive receivers (NA).
- **Depth observations.** These are defined in a `DataFrame`. Each row is a time step. Numbers define individual depths (m).

```
# Define study area
bathy = GeoArrays.read("../data/input/map-wahoo-100m.tif");
```

```
# Define timeline
start = DateTime(2025, 1, 1, 0, 0)
stop = DateTime(2025, 1, 31, 23, 58)
timeline = collect(start:Minute(2):stop)
first(timeline, 3)
```

```
3-element Vector{DateTime}:
 2025-01-01T00:00:00
 2025-01-01T00:02:00
 2025-01-01T00:04:00
```

```
# Define simulated path
path = CSV.read("../data/input/sim/example/path.csv",
                DataFrame)
first(path, 3)
```

```
3×6 DataFrame
 Row │ individual_id  timestamp            timestep  map_value  x          y   …
     │ Int64          String31             Int64     Float64    Float64    Flo …
─────┼──────────────────────────────────────────────────────────────────────────────
   1 │             1  2025-01-01                  1    50.1744  7.0895e5   6.2 …
   2 │             1  2025-01-01 00:02:00         2    59.4647  7.08803e5  6.2
   3 │             1  2025-01-01 00:04:00         3    42.1349  7.08782e5  6.2
                                                                 1 column omitted
```

```julia
# Define receiver positions
moorings_df = CSV.read("../data/input/sim/example/moorings-wahoo.csv",
                       DataFrame)
first(moorings_df, 3)
```

```
3×3 DataFrame
 Row │ receiver_id  receiver_x  receiver_y
     │ Int64        Float64     Float64
─────┼──────────────────────────────────────
   1 │           2   7.19404e5   6.2735e6
   2 │           3   7.06472e5   6.254e6
   3 │           4   7.09758e5   6.26773e6
```

```julia
# Define acoustic observations (0, 1)
acoustic_obs_df = CSV.read("../data/input/sim/example/acoustics-wahoo.csv",
                           DataFrame,
                           dateformat    = "yyyy-mm-dd H:M:S",
                           missingstring = "NA")
acoustic_obs_df[1:3, 1:10]
```

```
3×10 DataFrame
 Row │ 2      3      4      5      7      8      9      10     11     12
     │ Int64  Int64  Int64  Int64  Int64  Int64  Int64  Int64  Int64  Int64
─────┼──────────────────────────────────────────────────────────────────────
   1 │     0      0      0      0      0      0      0      0      0      0
   2 │     0      0      0      0      0      0      0      0      0      0
   3 │     0      0      0      0      0      0      0      0      0      0
```

```julia
# Define depth observations
depth_obs_df = CSV.read("../data/input/sim/example/depths-wahoo.csv",
                        DataFrame,
                        dateformat = "yyyy-mm-dd H:M:S")
first(depth_obs_df, 3)
```

```
3×1 DataFrame
 Row │ depth
     │ Float64
─────┼─────────
   1 │ 56.1585
   2 │ 59.2452
   3 │ 43.9744
```

#### Map

This is a map of the study area with the simulated path and receiver array added:

```
# Define bbox
ext = GeoArrays.bbox(bathy)
xmin, xmax = ext.X
ymin, ymax = ext.Y

# Plot bathymetry
Plots.plot(bathy,
           xticks = range(xmin, xmax, length = 3),
           yticks = range(ymin, ymax, length = 3))

# Add path
Plots.plot!(path.x, path.y;
            seriestype = :path, linewidth = 0.1,
            legend = false)

# Add receiver positions
Plots.scatter!(moorings_df.receiver_x, moorings_df.receiver_y)
```

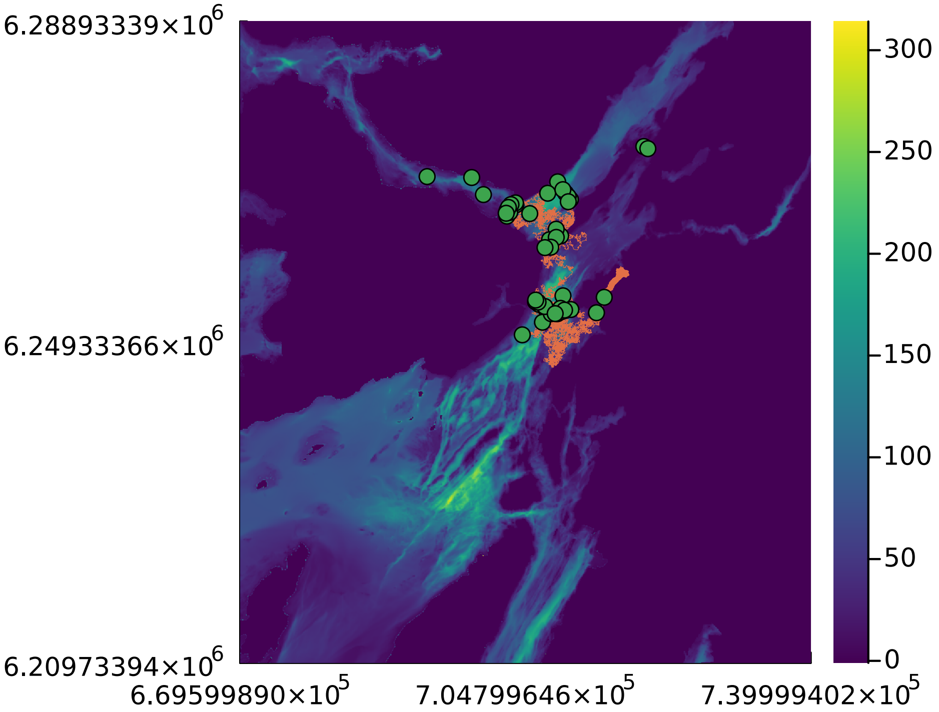


#### Model formulation

Now we formulate our state-space model. This comprises two sub-models: a movement model and a stack of observation models (one for each sensor).

#### Movement model

Wahoo.jl implements a Gaussian random walk (diffusion) movement model via two-dimensional convolution. We set the standard deviation of the movement model as movement_std. We select movement_std = 100 m based on previous research (Lavender et al., 2025). For this workflow, we treat movement_std (and the static parameters in the observation models) as known and focus inference on the latent states. (However, we note that in principle it is possible to estimate static parameters via the log-likelihood if data are sufficient and the computations can be managed.)

```
movement_std = 100.0;
```

#### Observation models

Observation models provide the link between the latent states and our observations. Wahoo.jl expects observations to be provided in a Vector, with one element for each sensor (e.g., receiver). Each element is a time series of observations for that sensor (with one observation per time step). For each sensor, a corresponding observation model should be provided. Wahoo.jl includes built-in observation models or we can define custom models with the signature:

```
function p_obs_*(signals, t::Int, depth::Number, distance::Number; kwargs...)
    _p_obs_*(signals[t], depth, distance; kwargs...)
end
```

##### Acoustic observation model

The acoustic observation model acts on acoustic observations. We provide acoustic observations in a Vector, with one element for each sensor. Each element is a time series of acoustic observations (0, 1 or −1) for that sensor. We use a Bernoulli observation model in which the probability of a detection is assumed to decline logistically with distance from a receiver. We use a model in which the rate of decline is controlled by two parameters, receiver_alpha and receiver_beta, informed by prior research (Lavender et al., 2025).

```
# Define a Vector with receiver positions:
acoustic_pos = tuple.(moorings_df.receiver_x, moorings_df.receiver_y);

# Collate acoustic observations in a Vector
# * Code observations as:
#   - 1 (detection)
#   - 0 (non-detection)
#   - -1 (no signal: inactive receiver)
# * We define a Vector of Vectors (each element is a sensor)
acoustic_obs = coalesce.(transpose(Array(acoustic_obs_df)), -1)
acoustic_obs = [acoustic_obs[i, :] for i in 1:size(acoustic_obs, 1)];

# Define acoustic observation model function
function p_obs_acoustic(signals, t::Int, waterdepth::Number, distance::Number;
                        receiver_alpha = 4.0, receiver_beta = -0.00940)
```

```julia
    signal = signals[t]
    if signal == 0
        return 1 - NNlib.sigmoid(receiver_alpha + receiver_beta * distance)
    end
    if signal == 1
        return NNlib.sigmoid(receiver_alpha + receiver_beta * distance)
    end
    return one(waterdepth)
end

# Define a Vector of observation models (one for each sensor)
p_obs_acoustic_vect = [p_obs_acoustic for i in 1:length(acoustic_obs)];
```

##### Depth observation model

As we have depth observations, we also define a depth observation model. Depth observations are collated in a `Vector`, as for acoustic observations. We define a truncated Gaussian observation model, with a mean on the seabed (in line with the benthic habit of skate) and a standard deviation that captures uncertainty in the bathymetric and depth measurements (based on prior research).

```julia
# Collect depth observations in a Vector
depth_obs = Vector(depth_obs_df.depth)

# Define depth observation model
function p_obs_depth(signals, t::Int, waterdepth::Number, distance::Number;
                     depth_std = 12.75270)
    signal = signals[t]
    if waterdepth < 0
        return zero(waterdepth)
    else
        mu = waterdepth
        f_obs = (1.0 / (sqrt(2.0 * pi) * depth_std)) *
                    exp(-((signal - mu)^2) / (2.0 * depth_std^2))
        lwr = (0.0 - mu) / (sqrt(2.0) * depth_std)
        upr = (350.0 - mu) / (sqrt(2.0) * depth_std)
        Z = 0.5 * (1.0 + erf(upr)) - 0.5 * (1.0 + erf(lwr))
        return f_obs / Z
    end
end;
```

#### Inference

Now we can run the inference routine. A single function, `track()`, is exported for this purpose. We provide `track()` with a probability distribution for the animal's initial location (coded as a `Matrix`). For simplicity, we select a uniform distribution over the marine area. We also supply our bathymetric raster and its spatial resolution, the time points at which we want to record probability distributions and our movement and observation models.

```julia
# Define probability distribution for the individual's initial location
p0 = deepcopy(bathy)
p0[p0 .>= 0] .= 1
p0[p0 .== -1] .= 0
p0 = p0 ./ sum(p0)
p0 = Matrix(p0);
```

```julia
# Define save time points as an _integer_
tsave = 1:60:length(timeline);
```

```julia
# Run inference
outfile = "../data/output/sim/example/Julia/wahoo.jld2"
if !isfile(outfile)

    println("Running Wahoo.jl...")

    res = track(pos_init            = p0,
                bathymetry          = bathy,
                spatial_resolution = 100,
                tsave               = tsave,
                movement_std        = movement_std,
                observations        = [depth_obs, acoustic_obs...],
                observation_models = [p_obs_depth, p_obs_acoustic_vect...],
                sensor_positions    = [nothing, acoustic_pos...],
                filter              = true,
                smoother            = true,
                n_trajectories      = 1,
                show_progressbar    = true,
                precision           = Float64)

    @save outfile res

else

    println("Loading Wahoo.jl results...")
    @load outfile res

end

  # Examine summary
  # * track() returns a NamedTuple:
  # * res.pos_filter
  #   > Array{Float, 4}: Ny × Nx × 1 × time of Prob(s_t | y_{1...t})
  # * res.pos_smoother
  #   > Array{Float, 4}: Ny × Nx × 1 × time of Prob(s_t | y_{1...T})
  # * res.residence_dist
```

```
  #   > Matrix{Float}: Ny × Nx of 1/T Sum Prob(s_t | y_{1...T})
  # * res.trajectories
  #   > Vector of trajectories from Prob(s_{1...T} | y_{1...T})
  # * res.log_p
  #   > Vector of Prob(y_t)
  # * res.tsave
  #   > Vector of time points
summary(res)
```

```
Loading Wahoo.jl results...
```

```
"@NamedTuple{log_p::Vector{Float64},      tsave::StepRange{Int64,      Int64},
trajectories::Vector{Matrix{Int64}},      pos_smoother::Array{Float64,      4},
residence_dist::Matrix{Float64}, pos_filter::Array{Float64, 4}}"
```

#### Analysis

You can analyse `Wahoo.jl`'s outputs using standard tools available in the `Julia` ecosystem or your preferred programming language. In this project, the above `Julia` code was integrated into a larger `RStudio Project`, so we collated the outputs in `R`. See the accompanying `R` workflow for details.
